# Cycling Molecular Assemblies for Selective Cancer Cell Golgi Disruption

**DOI:** 10.1101/2025.01.05.631374

**Authors:** Weiyi Tan, Qiuxin Zhang, Zhiyu Liu, Kangqiang Qiu, Divyanshu Mahajan, Thomas Gerton, Noah Copperman, Chaoshuang Xia, Cheng Lin, William Lau, Mikki Lee, Isabela Ashton-Rickardt, Pengyu Hong, Daniela Dinulescu, Jer-Tsong Hsieh, David Loeb, Ronny Drapkin, Jiajie Diao, Lei Lu, Bing Xu

**Affiliations:** Department of Chemistry, Brandeis University, Waltham, MA 02453, United States; Department of Cancer Biology, University of Cincinnati College of Medicine, Cincinnati, OH 45267, United States; School of Biological Sciences, Nanyang Technological University, Singapore 637551, Singapore; Department of Pathology, Brigham and Women’s Hospital, Harvard Medical School, Boston, MA 02115, United States; Department of Developmental & Molecular Biology, Albert Einstein College of Medicine, Bronx, NY 10461, United States; Department of Biochemistry & Cell Biology, Center for Biomedical Mass Spectrometry, Boston University Chobanian & Avedisian School of Medicine, Boston, MA 02118, United States; Department of Computer Science, Brandeis University, Waltham, MA 02453, United States; Department of Urology, Southwestern Medical Center, University of Texas, Dallas, TX 75235, United States; Ovarian Cancer Research Center, University of Pennsylvania, Perelman School of Medicine, Philadelphia, PA, 19104, United States

**Keywords:** Golgi, trafficking, enzyme switch, assemblies, cancer

## Abstract

The Golgi apparatus is a critical organelle responsible for intracellular trafficking and signaling, orchestrating essential processes such as protein and lipid sorting^1–5^. Dysregulation of its function has been implicated in various pathologies, including obesity, diabetes, and cancer, highlighting its importance as a potential therapeutic target. Despite this, the development of tools to selectively target the Golgi in specific cell types remain a significant unmet challenge in imaging and drug discovery. Golgi-specific enzyme activities, such as those mediated by protein acyltransferases and thioesterases^6^, offer an untapped opportunity to develop subcellularly localized therapeutics. Current approaches predominantly rely on direct protein binding but lack the necessary cell selectivity^7^, underscoring the unmet need for innovative strategies to selectively disrupt Golgi function in cancer cells. Here, we report the development of cycling molecular assemblies (CyMA), a novel class of small peptide derivatives (e.g., dipeptides), which exploit the unique enzymatic environment of the Golgi to establish futile cycles of reversible S-acylation. These assemblies selectively accumulate in cancer cell Golgi, interfering with protein S-acylation cycles and disrupting organelle homeostasis. CyMA impair key Golgi functions, including protein trafficking, glycosylation, and secretion, while demonstrating selective sparing hepatocytes and immune cells such as M1 macrophages. This selective activity represents a paradigm shift, utilizing an enzyme switch and leveraging intracellular environment rather than direct protein binding. Unlike conventional approaches, CyMA reduce tumor growth, drug resistance, and metastasis by pleiotropically disrupting Golgi related functions. By demonstrating the potential of futile cycles as a therapeutic strategy^8^, this study introduces a generalizable method for targeting organelle-specific enzyme activities. These findings not only underscore the therapeutic potential of CyMA in cancer but also pave the way for future applications in other Golgi-associated diseases.

## Main

The Golgi apparatus is a central trafficking and signaling hub in eukaryotic cells, regulating interorganellar communication^1,2,9^, lipid biosynthesis and trafficking^10,11^, cargo processing through glycosylation^12^ and lipidation^3^, and even cell death initiation^13^. Beyond its essential physiological roles, the Golgi plays critical roles in cancer proliferation and metastasis, as key proteins like IGF-1R^14^, ERBB2^15^, EGFR^16^, and TGFB1^17^ require Golgi-dependent glycosylation for functional integrity. Similarly, oncoproteins^18^ such as RAS^19^, MYC^20^, and WNT^21^ depend on Golgi-mediated glycosylation or lipidation, highlighting the organelle’s significance in cancer progression. Furthermore, the maturation of immune checkpoint proteins also requires Golgi functionality^22,23^, further highlighting its importance in oncogenic pathways. Despite the potential for Golgi-directed molecular imaging and therapy^24–26^, challenges remain, including limited cell selectivity, slow uptake, and mixed clinical outcomes^27^. Our recent findings show that thiophosphopeptides and peptide thioesters can rapidly target the Golgi, selectively killing cancer cells through enzyme-catalyzed, non-equilibrium self-assembly^28,29^. These insights prompt interest in refining Golgi-targeting strategies and further exploring the underlying mechanisms and applications. Here, we report a novel approach using cycling molecular assemblies (CyMA) that exploit a futile cycle established by counteracting acyltransferases and acyl-protein thioesterases (**Fig. 1a, b**). This dynamic process enable rapid accumulation within the Golgi at nanomolar concentrations via reversible acylation process mediated by palmitoyl acyltransferases (zDHHCs)^30,31^ and thioesterases (TEs: PPT1, LYPLA1, LYPLA2)^32^, which forms non-diffusive supramolecular assemblies that impede native protein palmitoylation, disrupt essential trafficking and posttranslational modifications (PTM), and downregulate phospho-AKT, thereby impairing RAS and RTK signaling. CyMA also induce organelle stress, destabilize the endoplasmic reticulum (ER), initiate autophagy, and increase ubiquitination of ribosomal proteins, culminating in selective cancer cell death without acquired resistance, while sparing normal and immune cells. *Ex vivo* studies on cancer spheroids and patient-derived tumor spheroids (PDOTS) and *in vivo* syngeneic murine models confirm CyMA’s potency. Moreover, CyMA modulate the tumor microenvironment by inhibiting CAF and M2 TAM, reducing pro-tumorigenic cytokines (e.g., TGFB1, VEGF, IL-10), and enhancing responses to immunotherapy. This previously unexplored strategy harnesses enzyme switches and reversible PTMs to achieve *in vivo* peptide lipidation with broad therapeutic implications.

**Figure 1.**
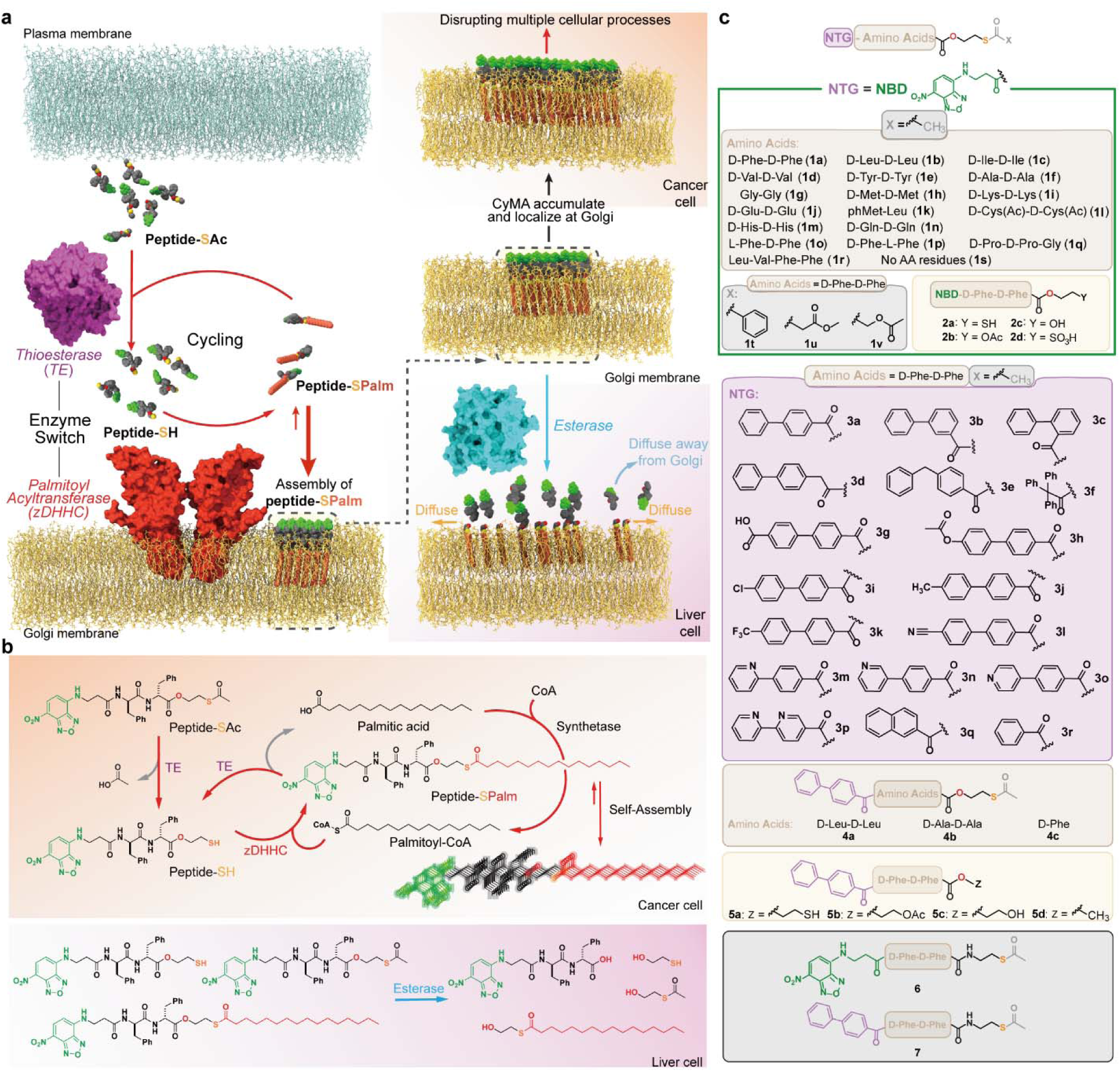
Selective accumulation of CyMA at the Golgi through an enzyme switch to establish futile cycles. (a) Schematic illustration of the futile cycle initiated by enzymatic deacylation of CyMA precursors, followed by selective accumulation at the Golgi through an enzyme switch involving thioesterases and palmitoyl acyltransferase. The consequences of CyMA accumulation at the Golgi in cancer and liver cells are illustrated. (b) Molecular structures and involved reaction pathways of CyMA precursors and metabolites within the futile cycle, along with exit from the cycle via hydrolysis of CyMA by esterases. (c) Molecular structures of CyMA precursors, relevant analogues and control compounds.

### Molecular design of CyMA

Our Golgi targeting thiophosphopeptides^28^ and peptide thioesters^29^ share several key features: (i) a nitrobenzoxadiazole (NBD) fluorophore that becomes highly fluorescent upon self-assembly^33^; (ii) a protease-resistant D-Phe-D-Phe dipeptide that promotes self-assembly^34^; and (iii) an enzyme-responsive trigger (thiophosphate or thioester) enabling enzyme-instructed self-assembly^35^ upon catalysis by alkaline phosphatases (ALP) or thioesterases (TEs). Although thiophosphopeptides are cell-selective based on ALP expression, their propensity for auto- dephosphorylation under acidic conditions limits broader applications. In contrast, peptide thioesters provide greater stability and Golgi localization but lack cell selectivity. To address these limitations, we introduced an ester linkage into peptide thioesters (**Fig. 1b, c**) to develop a stable, cell-selective Golgi-targeting platform. We designed a series of self-assembling peptides capped with NBD and incorporated ester and thioester bonds via 2-mercaptoethanol. This library includes dipeptides bearing a range of side chains: D-diphenylalanine (**1a**), D-dileucine (**1b**), D- diisoleucine (**1c**), D-divaline (**1d**), D-dityrosine (**1e**), D-dialanine (**1f**), diglycine (**1g**), D-dimethionine (**1h**), D-dilysine (**1i**), D-diglutamate (**1j**), L-photo-methionine-L-leucine (**1k**), D- di-S-acetylcysteine (**1l**), D-dihistidine (**1m**), and D-diglutamine (**1n**). To rule out receptor-specific interactions, we synthesized heterochiral precursors L-Phe-D-Phe (**1o**) and D-Phe-L-Phe (**1p**), as well as tripeptide (**1q**) and tetrapeptide (**1r**) motifs for generality. A non-self-assembling control (**1s**) was also prepared. Further modifying the acyl group (benzoyl (**1t**), 3-methoxy-3- oxopropanoyl (**1u**), acetoxyacetyl (**1v**)), and changing the thioester into free thiol (**2a**), acetyl ester (**2b**), hydroxyl (**2c**), and sulfonic acid (**2d**) allow us to examine the influence of these structural variations on Golgi targeting. Replacing NBD in the original thiophosphopeptide with a naphthyl group yielded a variant that potently inhibited HeLa cell proliferation (IC_50_ = 2.8 μM)^28^. Inspired by this result, we substituted NBD in **1a** with a more hydrophobic N-terminal biphenyl group, producing **3a**. Regioisomeric and heteroaromatic analogues (**3b–r**) were subsequently synthesized. We reduced the self-assembly propensity of **3a** by substituting its dipeptide with dileucine, dialanine, or phenylalanine (**4a–c**). Converting the acetyl thioester moiety in **3a** to free thiol, acetyl ester, or hydroxyl groups yielded **5a–c**, and employing a methyl ester (**5d**) would confirm that thioesters are necessary for in-cell CyMA formation. Finally, replacing the ester bond in **3a** with an amide linkage afforded canonical peptide thioesters **6** and **7** for underscoring the critical role of the ester bond in controlling CyMA dynamics.

### Golgi-targeting mechanism of CyMA

CyMA localize to the Golgi through three distinct steps (**Fig. 2a**): (I) TEs hydrolyze the thioester bond in CyMA precursors, producing thiol assemblies; (II) zDHHCs-mediated S-palmitoylation at the Golgi^3^ generates non-diffusive assemblies, while TE-catalyzed depalmitoylation sustains a dynamic cycle, resulting in strong green fluorescence from the NBD-labeled assemblies at the Golgi; (III) esterases hydrolyze the ester bond, producing hydrophilic molecules that subsequently disassemble and diffuse away.

**Figure 2.**
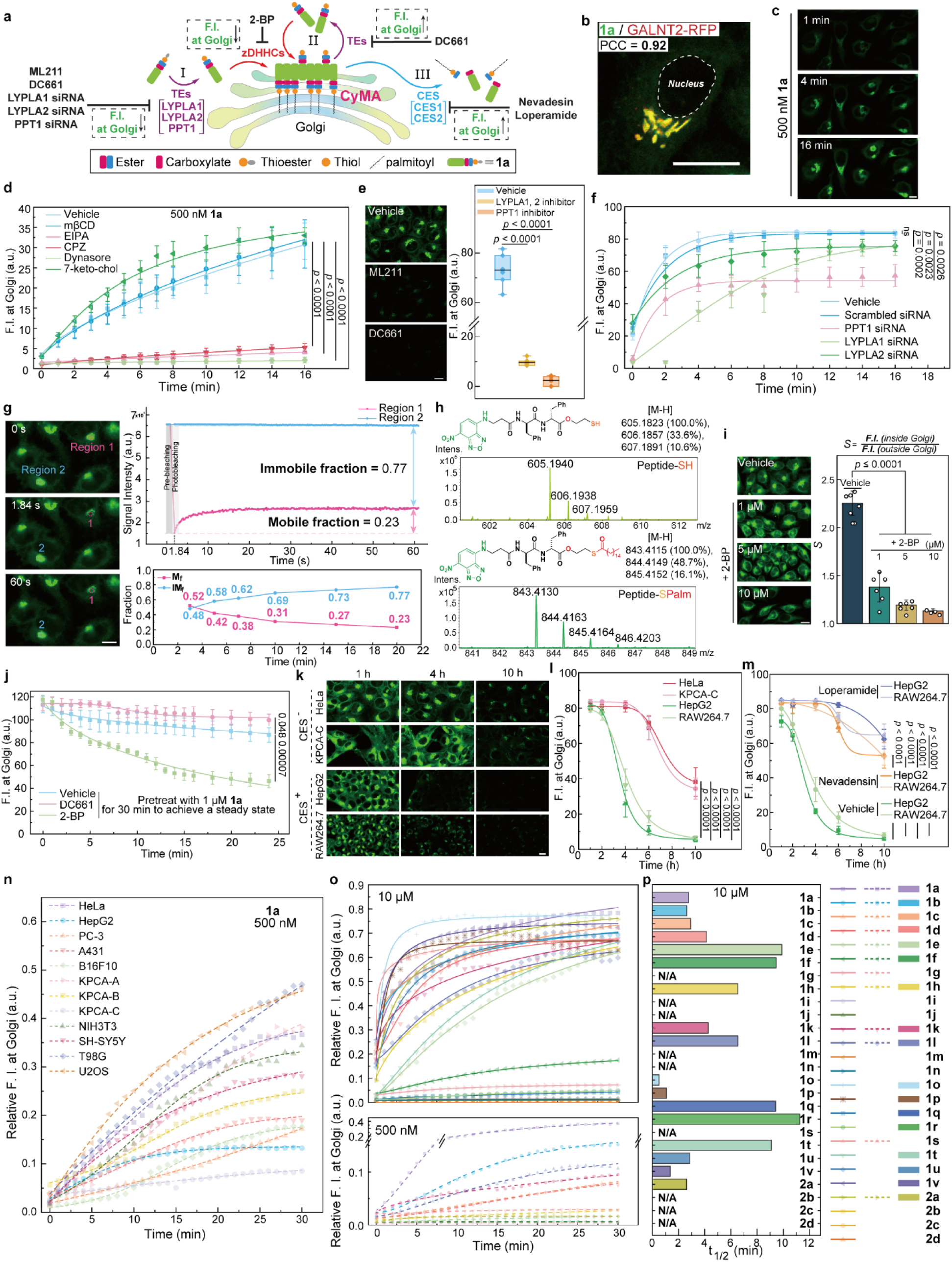
Golgi targeting features of CyMA precursors and the role of enzymes in futile cycle creation and disruption. (a) Schematic illustration of the intracellular enzymatic reactions involving CyMA precursors, showing the assembly and disassembly dynamics of CyMA at the Golgi. (i) CyMA precursors are hydrolyzed by TEs to form CyMA; (ii) zDHHCs palmitoylate the CyMA, resulting in selective accumulation at Golgi, where non-diffusive CyMA form and continuously cycle through palmitoylation-depalmitoylation; (iii) CyMA at the Golgi are hydrolyzed by esterases and dissociate from Golgi as small molecules and charged peptides. (b) CLSM of GALNT2-RFP HeLa cells treated with **1a** (1 μM, 10 min) and colocalization analysis. (c) CLSM of HeLa treated with **1a** (500 nM, 1, 4, 16 min). (d) Quantitative analysis of fluorescence intensity at the Golgi of HeLa pretreated with endocytosis inhibitors (mβCD, EIPA, CPZ, dynasore, 7-keto-chol) for 30 min, followed by treatment with **1a** (500 nM, 0 - 16 min) (n = 6). (e) CLSM and fluorescence intensity analysis at the Golgi of HeLa cells pretreated with vehicle, LYPLA1/2 inhibitor (ML211, 50 μM), or PPT1 inhibitor (DC661, 20 μM) for 30 min, then treated with **1a** (1 μM, 1h) (n = 6). (f) Quantitative analysis of the fluorescence intensity at the Golgi following treatment with **1a** (5 μM, 0 - 16 min) in wild-type HeLa and HeLa transfected with TE siRNA (PPT1, LYPLA1, LYPLA2 siRNA) (n = 5). (g) FRAP analysis of fluorescent Golgi after treatment with **1a** (2 μM, 20 min), with time-dependent mobile fraction (M_f_) and immobile fraction (IM_f_) at the Golgi following the treatment with **1a** (2 μM). (h) HRMS of the CyMA (**1a**) and the palmitoylated CyMA. (i) CLSM of HeLa cells pretreated with vehicle or DHHC inhibitor (2-BP, 1, 5, 10 μM) for 30 min and treated with **1a** (2 μM, 20 min), followed by “Golgi-specificity” analysis. (j) Quantitative analysis of the fluorescent intensity at the Golgi of HeLa cells pretreated with **1a** (1 μM, 30 min), then switched to fresh media with either PPT1 inhibitor (DC661, 20 μM) or zDHHC inhibitor (2-BP, 20 μM) (n = 6). (k) CLSM of CES negative (HeLa, KPCA-C) and CES positive (HepG2, RAW264.7) cells treated with **1a** (2 μM, 1, 4, 10 h). (l) Quantitative analysis of fluorescence intensity at the Golgi shown in (k) (n = 6). (m) Quantitative analysis of fluorescence intensity at the Golgi of CES positive (HepG2, RAW264.7) cells pretreated with vehicle, CES1 inhibitor (Nevadensin, 20 μM), or CES2 inhibitor (Loperamide, 20 μM) for 30 min, followed by treatment with **1a** (1 μM, 1, 2, 4, 6, 10 h) (n = 6). (n) Quantitative analysis of fluorescence intensity at the Golgi of various cell lines treated with **1a** (500 nM, 0 - 30 min) (n ≥ 6). (o) Quantitative analysis of fluorescence intensity at the Golgi of HeLa cells treated with various CyMA precursors at 10 μM or 500 nM (0 - 30 min) (n ≥ 6). (p) t_1/2_ of CyMA precursors with good Golgi targeting ability at 10 μM. Scale bar = 20 μm.

Fluorescence from CyMA precursor **1a** colocalized with Golgi with a Pearson correlation coefficient (PCC) of 0.92 (**Fig. 2b**), while showing minimal overlap with lysosomal, mitochondrial, or ER markers (**Supplementary Fig. S1a-c**). Golgi morphology became detectable within one minute of treatment with 500 nM of **1a** (**Fig. 2c**, **Supplementary Movie S1**), with fluorescence observed at the Golgi at concentration as low as 50 nM (**Supplementary Fig. S1d**, **Supplementary Movie S1**). Comparatively, the NBD-conjugated MGCTLSA peptide (10 μM), derived from the G sequence known to enable its Golgi localization^36^, showed minimal fluorescence at the Golgi (**Supplementary Fig. S1e-f**), highlighting CyMA’s specificity and efficacy. While **1a** colocalizes with GalT-RFP at 18 °C (**Supplementary Fig. S1g**), a temperature that specifically arrests the endosome-to-Golgi transport, no **1a** enters the cell at 0 °C, demonstrating that **1a** internalization requires active endocytosis. Endocytosis inhibitors, including chlorpromazine, EIPA, and or dynasore, significantly reduced Golgi fluorescence, while caveolin- and CLIC/GEEC-pathway inhibitors had negligible effect (**Fig. 2d**). These results indicate that CyMA internalize via clathrin- and dynamin-mediated endocytosis, macropinocytosis, and alternative pathways.

ML211 (a LYPLA1/2 inhibitor^37^) significantly reduced Golgi fluorescence, while DC661 (a PPT1 inhibitor^38^) blocked **1a** accumulation (**Fig. 2e**). Neither TE inhibitor substantially affected the Golgi accumulation of C6-NBD-Ceramide, and the Golgi structure remained intact following inhibitor pretreatment (**Supplementary Fig. S1h**). Knockdown of PPT1 and LYPLA1 reduced fluorescence accumulation rates, whereas LYPLA2 knockdown had a lesser effect (**Fig. 2f**, **Supplementary Fig. S1i**). These results confirm the essential role of TEs in CyMA activation. Fluorescence recovery after photobleaching (FRAP) experiment revealed a high immobile fraction (IM_f_ = 0.77) for CyMA assemblies at the Golgi, contrasting with the greater mobility of control C6-NBD-ceramide^24^ (IM_f_ = 0.22) (**Fig. 2g**; **Supplementary Fig. S1j**). These results indicate that CyMA’s high immobile fraction is attributed to its non-diffusive properties rather than Golgi compartmental stability^39^.

LC-HRMS analysis confirmed the lipidation of CyMA at the Golgi, identifying both thiol (Peptide-SH) and palmitoylated (Peptide-SPalm) forms of **1a** (**Fig. 2h**; **Supplementary Fig. S1k**). zDHHC inhibition by 2-BP^19^ resulted in a dose-dependent reduction of CyMA accumulation at the Golgi (**Fig. 2i**), with no detectable reaction between CyMA and 2-BP (**Supplementary Fig. S1l**)), emphasizing the importance of palmitoylation (i.e., switching from Peptide-SH to Peptide-SPalm) in Golgi targeting. 2-BP showed similar inhibition on CyMA with free thiol groups (**2a**) (**Supplementary Fig. S1m**), whereas 2-BP had no notable impact on C6- NBD-Ceramide (**Supplementary Fig. S1h**), further confirming that palmitoylation is essential for Golgi accumulation. Knockdown of several zDHHC isoforms slightly reduced fluorescence accumulation (**Supplementary Fig. S1n**), indicating the involvement of multiple zDHHC enzymes in this process.

To validate the palmitoylation-depalmitoylation cycle at the Golgi, cells were treated with TE (DC661) and zDHHC (2-BP) inhibitors after Golgi fluorescence had reached a steady state. Washing off **1a** from the media caused a gradual fading of Golgi fluorescence, indicating dynamic balance between Peptide-SPalm and Peptide-SH (**Fig. 2j**). TE inhibition increased fluorescence retention, while zDHHC inhibition diminished it. The varying effects of TE inhibition at different stages (**Fig. 2e**, **2j**) reflect the futile cycle at the Golgi. Early-stage TE inhibition halts the deacetylation of **1a** to Peptide-SH, preventing palmitoylation and thereby reducing fluorescence. At a later stage, TE inhibition maintains fluorescence by increasing the Peptide-SPalm/Peptide-SH ratio. Repeating these experiments with compound **6** and extended incubation (**Supplementary Fig. S2a-c**) confirmed that fluorescence changes were not due to esterase activity. A palmitoylation-independent Golgi probe, C6-NBD-ceramide, exhibited similar fluorescence fading under both inhibitors (**Supplementary Fig. S2d**). Extrinsically synthesized Peptide-SPalm, generated by substituting the acetyl group of **6** with a palmitoyl group (**Supplementary Fig. S2e**), showed limited cellular entry and formed extracellular aggregates due to its high hydrophobicity. These findings conclusively demonstrate the continuous palmitoylation-depalmitoylation cycles at the Golgi, with *in situ* palmitoylation being crucial for CyMA Golgi localization.

To examine influence of esterases on CyMA localization, cell lines with varying expression levels of carboxylesterase (CES) were examined. In HepG2 and RAW264.7 cells (high CES expression), Golgi fluorescence decreased significantly after 4 hours of **1a** treatment, while it remained stable in HeLa and KPCA-C cells (low CES expression) (**Fig. 2k**, **2l**). CES1 and CES2 inhibitors preserved Golgi fluorescence in HepG2 and RAW264.7 cells (**Fig. 2m**) but had minimal impact in HeLa cells (**Supplementary Fig. S3a**). Compound **6**, designed with an amide bond replacing the ester bond in **1a**, maintained Golgi fluorescence regardless of CES inhibition (**Supplementary Fig. S3b**), confirming that esterases modulate CyMA dynamics at the Golgi.

CyMA precursor **1a** functions as a general Golgi probe across various cell lines, with Golgi morphology becoming visible within 3 minutes of treatment with 500 nM **1a** (**Fig. 2n**; **Supplementary Fig. S3c**). Fluorescence accumulation varies across cell lines, likely due to differences in enzymatic activity and cellular environment. Replacing NBD fluorophore in **1a** with dansyl (DAN) or 4-(N,N-dimethylsulfamoyl)-2,1,3-benzoxadiazole (DBD) enabled imaging with different excitation wavelengths (**Supplementary Fig. S3d**), demonstrating CyMA’s versatility for imaging applications.

### Structural factors of CyMA’s Golgi-targeting

To elucidate the relationship between CyMA structure and Golgi localization, we systematically modified the molecular structure of precursor **1a** and evaluated their Golgi-targeting performance (**Fig. 2o** and **2p**). Substitution of D-diphenylalanine with D-dileucine (**1b**), D- diisoleucine (**1c**), or D-divaline (**1d**) resulted in slightly reduced fluorescence intensity at the Golgi under steady state condition. However, at 500 nM, fluorescence was markedly lower compared to **1a**, despite similar half-times (t_1/2_) to reach the steady state at 10 μM. This suggests that efficient self-assembly is crucial for Golgi targeting. CyMA precursors containing D- dityrosine (**1e**), D-dialanine (**1f**), diglycine (**1g**), and D-dimethionine (**1h**) exhibited reduced Golgi targeting ability. This reduction likely stems from their susceptibility to oxidization, attenuated self-assembly capacity, and lower hydrophobicity. While thioesters **1e** and **1f** targeted the Golgi at both 10 μM and 500 nM, their amide bond analogs failed to accumulate (**Supplementary Fig. S4a**), underscoring the necessity of a thiol group adjacent to an ester moiety for effective Golgi targeting. Charged CyMA precursors, such as D-dilysine (**1i**) and D-diglutamate (**1j**), showed minimal Golgi accumulation, highlighting the importance of hydrophobicity. Conversely, CyMA precursor with L-photo-methionine-L-leucine (**1k**) localized efficiently at the Golgi with t_1/2_ around 4 minutes (**Supplementary Fig. S4b**) and facilitated photo-crosslinking reactions^40^ at the Golgi. Proteomic analysis of photo-crosslinked proteins by **1k** revealed minimal Golgi-derived proteins (**Supplementary Fig. S4c**), suggesting that peptide-protein interactions are not the primary driver of Golgi localization.

Enhancing potential palmitoylation sites by incorporating D-diacetylcystine (**1l**) facilitated Golgi accumulation at both 10 μM or 500 nM (**Supplementary Fig. S4d**), though at a slower rate than **1a**. Precursors with D-dihistidine (**1m**) and D-diglutamine (**1n**) demonstrated good aqueous solubility due to protonation and hydrogen bond formation with water but failed to achieve robust self-assembly and Golgi targeting. Heterochiral precursors, including L-Phe-D-Phe (**1o**) or D-Phe-L-Phe (**1p**), exhibited Golgi targeting comparable to **1a**, thereby excluding receptor-mediated localization binding as a determining factor. Tripeptide (**1q**) and tetrapeptide (**1r**) precursors also targeted the Golgi, albeit less efficiently than **1a**, suggesting that peptide sequences modulate targeting efficiency. Notably, the control precursor **1s**, which lacks peptide segments, localized at the Golgi at a slower rate, emphasizing the generality of this thioester motif and the critical importance of self-assembly.

The role the acyl group in Golgi targeting was assessed by replacing the acetyl group of **1a** with benzoyl (**1t**), 3-methoxy-3-oxopropanoyl (**1u**), or acetoxyacetyl (**1v**). Among these analogs, **1t** exhibited the least effective Golgi targeting, likely due to the steric hinderance and conjugation effects associated with the aromatic ring, which impede enzymatic hydrolysis of the thioester bond. In contrast, **1u** showed a t_1/2_ similar to **1a**, while **1v** displayed a shorter t_1/2_, likely reflecting electronic effects. The lower pKa of the β-dicarbonyl in **1u** reduces the reactivity of its thioester bond under physiological conditions, compared to **1v**. Further, replacing the acetyl thioester of **1a** with a free thiol (**2a**), acetyl ester (**2b**), hydroxyl (**2c**), or sulfonic acid (**2d**) revealed the critical role of the thiol group in Golgi targeting. While **2a** displayed a similar t_1/2_ to **1a** at 10 μM, its Golgi fluorescence was significantly reduced at 500 nM, suggesting that free thiols engage off-target reaction prior to reaching the Golgi. None of the other modification (**2b-d**) accumulated at the Golgi, confirming that the thiol group is indispensable for enzymatic S-palmitoylation and efficient Golgi targeting.

### Cell-selective cytotoxicity of CyMA in vitro and in vivo

CyMA compounds display cell-selective activity by leveraging unique cycling mechanisms at the Golgi and eliciting cellular responses that vary by cell type. At low precursor concentrations, they induce cell death and inhibit migration in various cancer cell lines (**Fig. 3a**). This anticancer efficacy extends to three-dimensional (3D) cancer models, including cancer cell spheroids, PDOTS, and syngeneic murine tumor models. Furthermore, cells treated with CyMA exhibit minimal acquired drug resistance. Notably, CyMA precursors synergize with immune checkpoint blockade (ICB), highlighting their potential both as a monotherapy and in combination with immunotherapy. In cells overexpressing CES, CyMA undergo hydrolysis and diffuse from the Golgi, rendering them non-cytotoxic, which is further evident by restoring CyMA’s cytotoxicity in CES-overexpressing cells through inhibiting CES by bis(p-nitrophenyl) phosphate (BNPP) (**Fig. 3a**).

**Figure 3.**
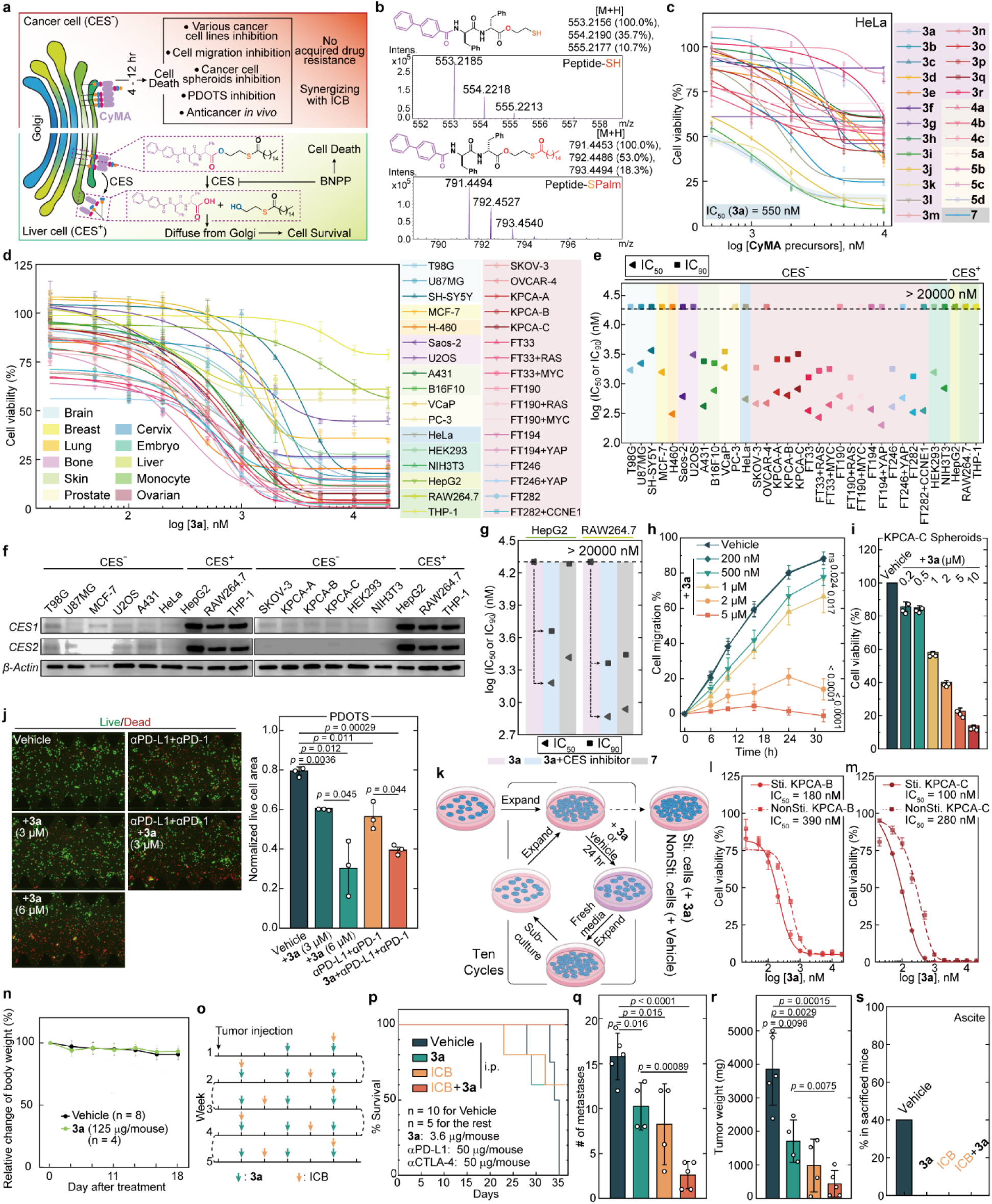
Cell-selective antitumor activity of CyMA *in vitro*, *ex vivo*, and *in vivo*. (a) Schematic illustration showing the cell-selective feature of CyMA precursors due to varying CES expression levels in cells. (b) HRMS analysis of deacylated and palmitoylated CyMA (**3a**) in cell lysate. (c) Cell viability of HeLa cells treated with CyMA precursors for 24 hours (n = 3). Cell viability of various cell lines treated with the CyMA precursor, **3a**, (n = 3). (e) IC_50_ and IC_90_ values of **3a** against the cell lines shown in (d). (f) Immunoblotting of CES1 and CES2 expression in representative cell lines from (d). (g) IC_50_ and IC_90_ values of **3a** and **7** against CES overexpressed cell lines (HepG2, RAW264.7) co-incubated with or without CES inhibitor (BNPP, 50 μM) for 24 hours. (h) Percentage of cell migration of HeLa cells treated with **3a** (n = 3). (i) Cell viability of KPCA-C cell spheroids treated with **3a** for 72 hours (n = 3). (j) Live/dead cell staining of PDOTS treated with **3a** combined with or without immune checkpoint inhibitors (200 μg/mL anti-PD-1 and anti-PD-L1) (n = 3). (k) Schematic illustration of CyMA-stimulated cells (Sti. cells) versus non-stimulated cells (NonSti. cells). (l) Cell viability of stimulated or non-stimulated KPCA-B cells treated with CyMA precursors (**3a**) for 24 hours (n = 3). (m) Cell viability of stimulated or non-stimulated KPCA-C cells treated with CyMA precursors (**3a**) for 24 hours (n = 3). (n) Relative change in body weight of mice injected with vehicle or **3a** (125 μg/mouse, three times a week for total 8 injections) intraperitoneally in PBS. (o) Workflow of the *in vivo* study combining **3a** with or without ICB. (p) Survival curves, (q) Number of metastases, (r) Tumor weight and (s) presence of ascites in KPCA-B bearing mice treated with vehicle, **3a** alone, ICB alone, or the combination of **3a** and ICB (n = 5 for vehicle or the combination, while n = 4 for **3a** or ICB).

To refine the structural characteristics of CyMA, we substituted the NBD fluorophore in **1a** with various N-terminal groups (NTGs), generating CyMA precursors **3a**-**3r**. They retained the Golgi-targeting thioester warhead that undergoes continuous deacylation and palmitoylation cycles. The para-biphenyl motif in **3a** proved critical, as **3a** was deacylated and palmitoylated in cells (**Fig. 3b**), displayed pronounced Golgi localization (**Supplementary Fig. S5a**) and potent cytotoxicity (IC_50_ ∼ 550 nM, **Fig. 3c**) that was much lower than **1a** (**Supplementary Fig. S5b**). Structural variations—such as meta-/ortho-biphenyl (**3b**, **3c**), altered biphenyl linkers (**3d**, **3e**), and triphenylmethyl substitution (**3f**)—reduced cytotoxicity, suggesting that **3a**’s unique supramolecular packing is integral to its efficacy. Induction of carboxyl or acetate groups (**3g**, **3h**) lowered efficacy, likely due to increased hydrophilicity. Conversely, para-substitutions (**3i**-**3l**) generally preserved cytotoxicity, indicating a self-assembly-driven mechanism rather than conventional ligand-receptor interactions. Heterocyclic NTGs (**3m**-**3p**) yielded milder cytotoxicity, owing to enhanced aqueous solubility that disfavors self-assembly. Naphthoyl (**3q**) and phenyl (**3r**) groups were less effective than biphenyl. Similarly, replacing the diphenylalanine motif with dileucine (**4a**), dialanine (**4b**) or phenylalanine (**4c**) significantly reduced cytotoxicity, reinforcing the importance of self-assembly motifs. Additional modifications—replacing the thioester in **3a** with a thiol (**5a**), ester (**5b**), hydroxyl (**5c**), or methyl ester (**5d**)—lowered the efficacy, likely because these groups fail to participate in the palmitoylation-depalmitoylation cycle *in situ* at Golgi. Amide bond substitution (**7**) also decreased cytotoxicity, indicating that ester-containing peptide assemblies are more effective in eliciting cellular responses than canonical peptide assemblies. We selected **3a** as the representative CyMA precursor due to its potency and low molecular weight, and we more extensively assessed its cytotoxicity and anticancer activity both *in vitro* and *in vivo*.

Testing **3a** across a broad panel of human and murine cell lines—including those derived from brain (T98G, U87MG, SH-SY5Y), breast (MCF-7), lung (H-460), bone (Saos-2, U2OS), skin /melanoma (A431, B16F10), prostate (VCaP, PC-3), cervix (HeLa) cancer cell lines, as well as embryonic-derived cell lines (HEK293, NIH3T3), hepatocytes (HepG2), monocytes (RAW264.7, THP-1), and ovarian cancer cell lines (SKOV-3, OVCAR-4, KPCA-A, KPCA-B, KPCA-C, FT33, FT190, FT194, FT246, FT282, and derivatives)—revealed pronounced cytotoxicity against most ovarian cancer cell lines and certain other lines, including H460, Saos-2, A431, B16F10, and HeLa cells (**Fig. 3d**), with IC_50_ values around 500 nM after 24 h (**Fig. 3e**). Notably, the IC_90_ values of **3a** for A431, B16F10, SKOV-3, KPCA-A, KPCA-B, KPCA-C, FT33, FT33+RAS, FT33+MYC, FT190+RAS are below 2 μM (1.2 μg/mL) (**Fig. 3e**), underscoring CyMA’s potent anticancer efficiency and translational potential. Interestingly, while PC-3 cells exhibited resistance at 24 h, sensitivity increased after 72 h (IC_50_ ∼ 600 nM, **Supplementary Fig. S5c**), indicating variability in temporal response. In contrast, cells overexpressing CES (HepG2, RAW264.7, THP-1; **Fig. 3f**) remained viable (IC_50_ > 20 μM) (**Fig. 3e**), demonstrating selective cytotoxicity (**Fig. 3d-f**). Replacing the ester in **3a** with an amide bond (**7**) resulted in uniformly inhibited cell growth regardless of CES expression (**Supplementary Fig. S5d**), whereas CES inhibition by BNPP drastically enhanced **3a**’s cytotoxicity in HepG2 and RAW264.7 cells, aligning these values with those of compound **7** (**Fig. 3g**).

CyMA inhibited cell migration in a concentration-dependent manner, with partial inhibition at near-IC_50_ concentrations and complete inhibition at higher doses (**Fig. 3h**; **Supplementary Fig. S5e**). In 3D tumor spheroid models, **3a** (5 μM) reduced KPCA-C spheroid viability by ∼80% (**Fig. 3i**). Consistent with this, PDOTS showed marked, concentration-dependent reductions in live cells upon treatment (**Fig. 3j**). Since PDOTS retain essential features of the native tumor immune microenvironment^41^, we combined CyMA with immune checkpoint blockade (ICB; anti-PD-L1 and anti-PD-1) and observed enhanced the proliferation inhibition (**Fig. 3j**), suggesting a promising synergistic effect. To assess the potential for minimizing acquired drug resistance, we stimulated KPCA-B and KPCA-C cells with **3a** and assessed their subsequent response (**Fig. 3k**). Strikingly, IC_50_ values that were initially 390 nM and 280 nM for KPCA-B and KPCA-C cells, respectively, decreased to 180 nM and 100 nM after stimulation (**Fig. 3l**, **3m**). These results indicate that, unlike conventional inhibitors, CyMA not only circumvents resistance mechanisms but also potentiates cellular responsiveness to treatment.

Finally, we evaluated the *in vivo* antitumoral efficacy in a syngeneic tumor model, with a particular focus on its combination with immunotherapy, as Golgi regulate the trafficking of immunosuppressive cytokines (*vide infra,* **Fig. 6**). Intraperitoneal injection of **3a** (125 μg/mouse, three times a week for eight doses) were well-tolerated (**Fig. 3n**). In KPCA-B-bearing C57BL/6 mice, a low dose of **3a** (i.p.; 3.6 μg/mouse, 0.144 mg/kg) significantly improved survival rates compared to vehicle-treated controls, whereas much higher doses of ICB antibodies (50 μg/mouse each for both anti-PD-L1 and anti-CTLA-4) were required to achieve similar survival benefits (**Fig. 3o, p**). Both CyMA and ICB reduced metastases, tumor weight, and the incidence of ascites (**Fig. 3q-s**). The combination of CyMA and ICB led to a complete final survival (**Fig. 3p**) and reduced metastases (**Fig. 3q**), and further decreased tumor burden (**Fig. 3r**), consistent with observations from PDOTS and underscoring the strong synergism between CyMA and ICB.

Collectively, these findings highlight the necessity of further investigation into the cellular and molecular mechanisms through which CyMA induces cell death and modulates the TME to enhance ICB efficacy.

### Disruption of Golgi-related trafficking by CyMA

CyMA accumulate at the Golgi and eventually reach a steady state, thereby disrupting both anterograde and retrograde Golgi trafficking (**Fig. 4a**). Unlike golgin-mediated disruption of trafficking that relies on decorating target organelles with specific golgins^42^, CyMA accumulation at the Golgi membrane exerts a global blockade, affecting transport in both directions. Analysis of Golgi specific proteins revealed significant dispersion of KDELR2 and scattering of medial- and trans-Golgi proteins following CyMA treatment (**Fig. 4b**). Notably, even a 1-hour treatment with **3a** (5 μM) resulted in extensive Golgi fragmentation, as visualized with **1a** (**Supplementary Fig. S5f**), demonstrating that CyMA disrupt the entire Golgi network. Although Golgi can remain functional despite fragmentation in cancer cells, CyMA-mediated Golgi disruption significantly impedes Golgi functions.

**Figure 4.**
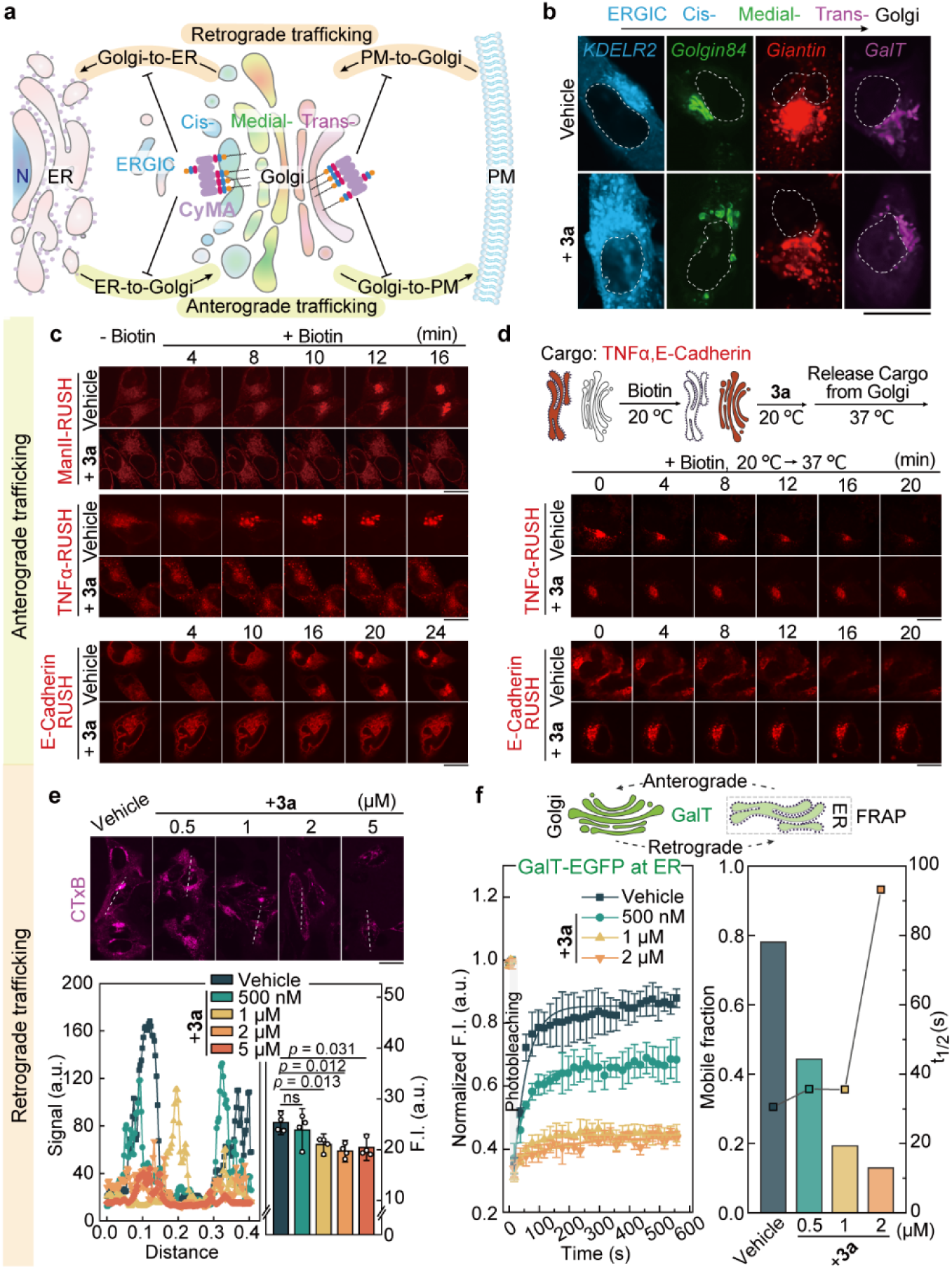
CyMA at Golgi disrupt intracellular trafficking. (a) Schematic illustration of Golgi disruption and intracellular trafficking blockade as the primary cellular responses triggered by CyMA. (b) CLSM images of ECFP-KDELR2, EGFP-Golgin84, mScarlet-Giantin and miRFP670-GalT in HeLa cells treated with or without **3a** (1 μM, 6 hours). (c) Time-lapse CLSM of ManII-mCherry-RUSH, TNFα-mCherry-RUSH, and E-Cadherin-mCherry-RUSH in HeLa cells treated with or without **3a** (2 μM, 6 hours), followed by biotin (40 μM) treatment. (d) Schematic illustration of Golgi-to-PM trafficking determination using the RUSH system; and time-lapse CLSM of TNFα-mCherry-RUSH and E-Cadherin-mCherry-RUSH in HeLa cells treated with or without **3a** (2 μM, 6 hours) at 20, followed by monitoring at 37. (e) CLSM of CTxB-AF647 in HeLa cells pretreated with or without **3a** at various concentrations for 6 hours, then treated with CTxB-AF647 (1 μg/mL, 1 hour) (n = 4). (f) FRAP analysis of the ER pool of GalT-EGFP with or without the treatment of **3a** at different concentrations for 6 hours, showing the quantified mobile fraction and t_1/2_. Scale bar = 20 μm.

To assess the impact on anterograde trafficking, we employed the Retention Using Selective Hooks (RUSH) system^43^ to track the transport of mannosidase II (ManII), TNF-α and E-Cadherin from the ER to the Golgi and onward to the plasma membrane (PM) (**Fig. 4c**, **4d**). In control cells, these proteins reached the Golgi within 10 min of biotin addition, whereas in CyMA-treated cells, the proteins were retained in the ER with no detectable Golgi localization (**Fig. 4c**). Unlike brefeldin A (BFA), which induces Golgi disassembly and redistribution into the ER^44^, CyMA uniquely disrupt Golgi morphology and function. To examine Golgi-to-PM export, we accumulated TNF-α and E-Cadherin in the Golgi at 20 °C in the presence of biotin, then shifted to 37 °C to initiate their transit to PM^43^. In untreated cells, Golgi fluorescence decreased, indicating successful trafficking to the PM, whereas in CyMA-treated cells no fluorescence decrease occurred, demonstrating a blockade in Golgi-to-PM transport (**Fig. 4d**).

CyMA also block retrograde trafficking. Using fluorophore-labeled cholera toxin subunit B (CTxB), a standard retrograde tracer^45^, we found that while control cells displayed Golgi-localized CTxB, CyMA-treated cells showed dispersed intracellular puncta with reduced total fluorescence intensity (**Fig. 4e**), indicating a severe impairment in PM-to-Golgi transport. To probe Golgi-to-ER retrograde transport, we performed FRAP on the ER pool of B4GALT1 (GalT), a protein that shuttles between the ER and Golgi^46^. Upon **3a** treatment, there was a concentration-dependent inhibition of GalT retrograde transport, evidenced by a reduced mobile fraction and an increased t_1/2_ (**Fig. 4f**, **Supplementary Fig. S6**). Collectively, these results show that CyMA potently disrupt Golgi organization and function by blocking both anterograde and retrograde trafficking, ultimately compromising the entire secretory pathway.

### Mechanisms of cancer cells inhibition by CyMA

As shown in **Fig. 5a**, CyMA treatment compromises Golgi integrity and disrupts intracellular trafficking, which subsequently impairs ER homeostasis and promotes mitochondrial fission. This disruption of trafficking and PTMs (e.g., lipidation and glycosylation) leads to protein mislocalization and dysfunction within critical signaling pathways, ultimately culminating in cell death. CyMA also induce autophagosome formation but paradoxically inhibit its maturation and disrupt basal polyubiquitination, further accelerating cell death. These findings suggest that CyMA exert cytotoxic effects through multiple pathways linked to trafficking disruption, thereby reducing the likelihood drug resistance.

**Figure 5.**
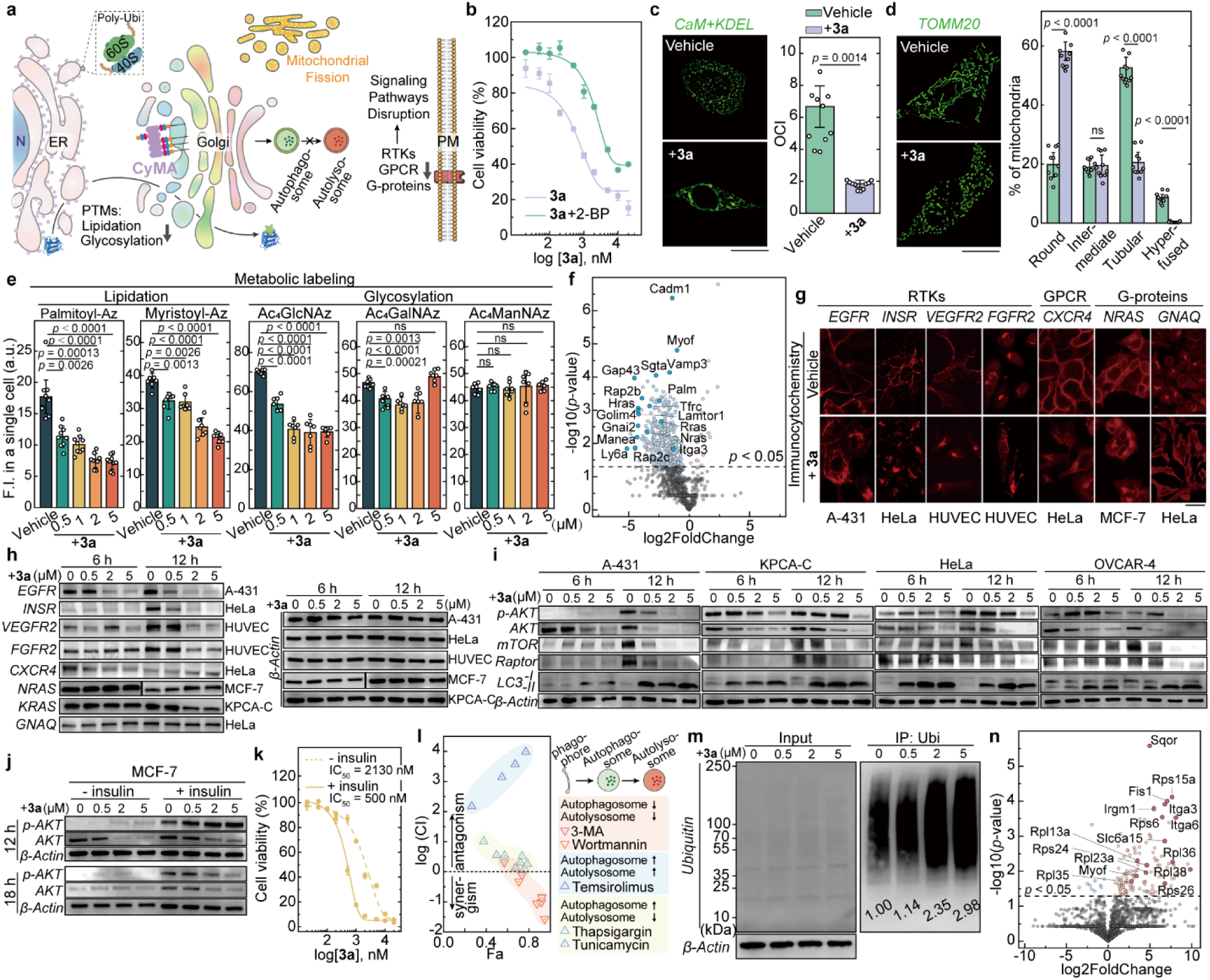
CyMA at Golgi lead to cell death by disrupting multiple cellular processes. (a) Schematic illustration of the secondary and indirect cellular responses triggered by CyMA. (b) Cell viability of HeLa cells treated with **3a**, with or without 2-BP (10 μM), for 24 hours (n = 3). (c) Structured illumination microscopy (SIM) images of the ER in HeLa cells treated with or without **3a** (2 μM, 24 hours) (n = 10). (d) SIM images of mitochondria in HeLa cells treated with or without **3a** (2 μM, 24 hours) (n = 10). (e) Analysis of protein lipidation and glycosylation in HeLa cells treated with **3a** at different concentrations for 12 hours via metabolic labeling (n = 10 for palmitoylation analysis, and n = 7 for the rest). (f) Volcano plot of the regulated palmitoylated proteins in KPCA-C cells treated with **3a** (2 μM, 6 hours) identified by LC-MS/MS (n = 3). (g) Immunocytochemistry of representative RTKs (EGFR, INSR, VEGFR2, FGFR2), GPCR (CXCR4) and G-proteins (NRAS, GNAQ) in different cell lines with or without the treatment of **3a** (2 μM, 6 hours). (h) Immunoblotting of the representative RTKs (EGFR, INSR, VEGFR2, FGFR2), GPCR (CXCR4) and G-proteins (NRAS, KRAS, GNAQ) in different cell lines with or without the treatment of **3a** (500 nM, 2 μM, 5 μM; 6 and 12 hours). (i) Immunoblotting of significant proteins (p-AKT, AKT, mTOR, Raptor, LC3B) in the AKT-mTOR signaling pathway in various cell lines with or without the treatment of **3a** (500 nM, 2 μM, 5 μM; 6 and 12 hours). (j) Immunoblotting of p-AKT and AKT in MCF-7 cells stimulated with or without insulin and treated with or without **3a** (500 nM, 2 μM, 5 μM; 12 and 18 hours). (k) Cell viability of insulin-stimulated and unstimulated MCF-7 cells treated with or without **3a** for 24 hours (n = 3). (l) Quantitative analysis of synergism and antagonism in combinations of **3a** with autophagy regulators; 3-methyladenine (3-MA) and Wortmannin inhibit autophagosome formation and further maturation; Temsirolimus activates autophagosome formation and autolysosome maturation; Thapsigargin and Tunicamycin activate autophagosome formation but block the autolysosome formation (n = 3). (m) Immunoprecipitation of ubiquitinated proteins in the lysates of KPCA-C cells treated with or without **3a** (500 nM, 2 μM, 5 μM, 6 hours), with quantitative analysis normalized over β-Actin. (n) Volcano plot of the ubiquitinated proteins in KPCA-C cells treated with **3a** (2 μM, 6 hours) identified by LC-MS/MS (n = 3). Scale bar = 20 μm.

Notably, the severe cell death induced by CyMA, particularly by compound **3a**, can be mitigated by inhibiting zDHHCs with 2-BP (**Fig. 5b**, **Supplementary Fig. S7a**). Knockdown of several zDHHC family proteins similarly reduces **3a**-mediated cytotoxicity (**Supplementary Fig. S7b**), indicating that palmitoylation, mediated by multiple zDHHCs, drives CyMA accumulation at Golgi and contributes to cell death. Beyond Golgi, CyMA also affect other organelles: following **3a** treatment, the ER becomes increasingly entangled (**Fig. 5c**, **Supplementary Fig. S7c**), and mitochondria adopt a rounded morphology indicative of fission (**Fig. 5d**, **Supplementary Fig. S7d**). By disrupting intracellular trafficking, CyMA impair protein lipidation and glycosylation. Metabolic labeling^47–49^ revealed a concentration-dependent reduction in palmitoylated and myristoylated proteins (**Fig. 5e**), accompanied by a significant decline in global glycosylation (**Supplementary Fig. S8a**), particularly O-GlcNAcylation, a key regulator of tumor progression^50^. O-GalNAcylation was modestly reduced at lower concentrations (i.e., 500 nM, 1 μM and 2 μM) but remained near basal levels at higher concentrations (5 μM), while sialylation remained unaffected (**Fig. 5e**). LC-MS/MS^51^ analysis of the palmitoyl proteome in **3a**-treated KPCA-C cells identified reduced palmitoylation (**Fig. 5f**; **Supplementary Table S1)**, including decreasing in Ras proteins (HRas, NRas, RRas), likely impairing their oncogenic functions^52^. Palmitoylation of Lamtor1, which regulate mTORC1 signaling and cell death^53^, was similarly affected, suggesting that the reduced palmitoylation of diverse proteins contributes to the cytotoxic effect of CyMA (**Fig. 5f)**.

We further explored CyMA’s effects on protein localization, including RTKs (EGFR, INSR, VEGFR2, FGFR2), a GPCR (CXCR4), and G-proteins (RAS, GNAQ) (**Fig. 5f**). Following **3a** treatment, EGFR, INSR and VEGFR2 relocated from the PM to the perinuclear region, while FGFR2, normally resident at the Golgi, dispersed into the cytoplasm. Unlike ligand- or inhibitor-induced internalization^54^, this mislocalization is attributed to trafficking disruption. For example, syntaxin-6, a SNARE-associated protein critical for RTK translocation at the Golgi^55,56^, is disrupted by CyMA (**Supplementary Fig. S8b**). Similarly, CXCR4^57^ and RAS^58^ relocated to the perinuclear region, and GNAQ no longer localized to the Golgi. Immunoblotting on the same cell lines revealed a significant reduction in the levels of four RTKs (EGFR, INSR, VEGFR2, FGFR2) (**Fig. 5h**). In contrast, CXCR4 levels transiently declined and then slightly recovered, NRAS remained stable in MCF-7 cells, KRAS was downregulated in KPCA-C cells, and GNAQ levels were unchanged. These results demonstrate that CyMA not only perturb the subcellular localization of key proteins but also alter their expression levels.

RTKs play central roles in cancer pathogenesis^59^, partly through activating the AKT pathway, which regulates growth, survival, and proliferation^60^. CyMA consistently suppressed AKT expression and phosphorylation, and reduced mTOR and Raptor expression across several cancer cell lines (A-431, KPCA-C, HeLa, OVCAR-4) (**Fig. 5i**) and in NIH3T3 embryonic fibroblasts (**Supplementary Fig. S8c**), indicating dampened mTORC1 activity. CyMA also modulated the Ras/Raf/MAPK pathway, evidenced by varied phosphorylation of ERK, P38, and JNK across cell lines (**Supplementary Fig. S8d**). An excitation-ratiometric assay^61^ confirmed reduced AKT kinase activity following CyMA treatment (**Supplementary Fig. S8e**). Enhanced CyMA sensitivity in insulin-stimulated MCF-7 cells^62,63^ (IC_50_ ∼ 500 nM) compared to unstimulated cells (IC_50_ ∼ 2.1 µM) further validated AKT pathway disruption as a mechanism of growth inhibition (**Fig. 5j, k**; **Supplementary Fig. S8f**). Although a 12-hour treatment did not markedly affect phosphorylated AKT levels, an 18-hour exposure reduced phospho-AKT in insulin-stimulated cells. These findings indicate that AKT pathway activation may enhance CyMA sensitivity, confirming that disruption of RTK-AKT signaling is a key mechanisms of CyMA-induced cell death.

CyMA-induced cell death involves additional pathways. Elevated LC3-II/LC3-I ratio and immunocytochemistry data (**Fig. 5i**; **Supplementary Fig. S9a**) indicate enhanced autophagosome formation^64^. However, an RFP-GFP tandem fluorescent-tagged LC3 assay^65^ suggested that CyMA block autophagosome maturation (**Supplementary Fig. S9b**). Co-incubating cells with autophagy regulators and subsequent CompuSyn^66^ analysis revealed that CyMA synergize with autophagy inhibitors (3-MA and wortmannin)^67^ but exhibit antagonism with the autophagy activator (temsirolimus^68^) or treatments promoting autophagosome formation without maturation (thapsigargin and tunicamycin)^69^ (**Fig. 5l**). Thus, while initial autophagosome formation may represent a stress response^70^, CyMA-induced inhibition of autophagosome maturation leads to cell death. Consistent with maturation arrest of the autophagosome, minimal mitophagy or ER-phagy can be observed (**Fig. 5c**, **5d**, **Supplementary Fig. S9c**,**d**). PI4KIIα dispersed into small cytoplasmic puncta after **3a** treatment (5 μ M) (**Supplementary Fig. S9e**) due to disrupted palmitoylation^71–73^, which is essential for autophagosome-lysosome fusion. A more condensed pattern of LC3 induced by CyMA, compared to chloroquine, suggested a unique mechanism of autolysosome inhibition (**Supplementary Fig. S9a**). These findings underscore the intricate role of autophagy in CyMA- induced cell death^74^.

In addition to autophagy dysregulation, CyMA affect the ubiquitin-proteasome system. A 6-hour treatment with 5 μM **3a** resulted in a greater than threefold increase in polyubiquitinated protein levels in KPCA-C (**Fig. 5m**) and HeLa cells (**Supplementary Fig. S10a**). LC-MS/MS analysis of polyubiquitinated proteins revealed increased polyubiquitination of several ribosomal proteins (**Fig. 5n**, **Supplementary Table S2**), suggesting the activation of stress-induced quality control mechanisms^75^. Proteins requiring palmitoylation (e.g., Itga3, Itga6, and Myof)^76–78^ also displayed elevated polyubiquitination, correlating with reduced palmitoylation levels (**Fig. 5f**). Elevated polyubiquitination of Sqor suggests a role for ferroptosis^79^, supported by RNA-Seq data showing upregulation of ferroptosis-related proteins (e.g., Gclm^80^, Hspb1^81^, Sqle^82^, Fads2^83^) (**Supplementary Fig. S10b**; **Supplementary Table S3**). Despite these insights, cell death outcomes are complex^84^ and depend on cell type and treatment duration. No canonical cell death inhibitors fully rescued the CyMA-treated cells (**Supplementary Fig. S10c**), highlighting the multifaceted nature of CyMA’s mechanisms of action.

### TME modulation by CyMA

TME is a dynamic ecosystem that fosters tumor progression, angiogenesis, metastasis, and immunoresistance, thereby posing challenges to cancer therapy^85^. Within the TME, cancer-associated fibroblasts (CAFs)—a major TME component—secrete cytokines that drive tumor growth and metastasis, rendering them a key hallmark of cancer^86^. Another critical TME constituent is the tumor-associated macrophage (TAM), which can exist in an antitumorigenic M1 state and or a pro-tumorigenic M2 state, with the latter serving as a promising therapeutic target^87^. Senescent tumor cells (STCs) also play pivotal roles in cancer progression by promoting collective invasion and establishing a cytokine barrier that protects non-senescent tumor cells from immune surveillance^88^. As shown in **Fig. 6a**, we demonstrated that CyMA effectively inhibit cancer cells, CAFs, and STCs, while selectively targeting M2 macrophages and sparing M1 macrophages, thus mitigating tumor-promoting cell populations with minimal toxicity to their antitumoral counterparts. Moreover, CyMA disrupt intracellular anterograde trafficking, impeding the secretion of key cytokines, including TGF-β1 and VEGF, which are released by CAFs, STCs, and cancer cells to sustain cancer progression and angiogenesis.

**Figure 6.**
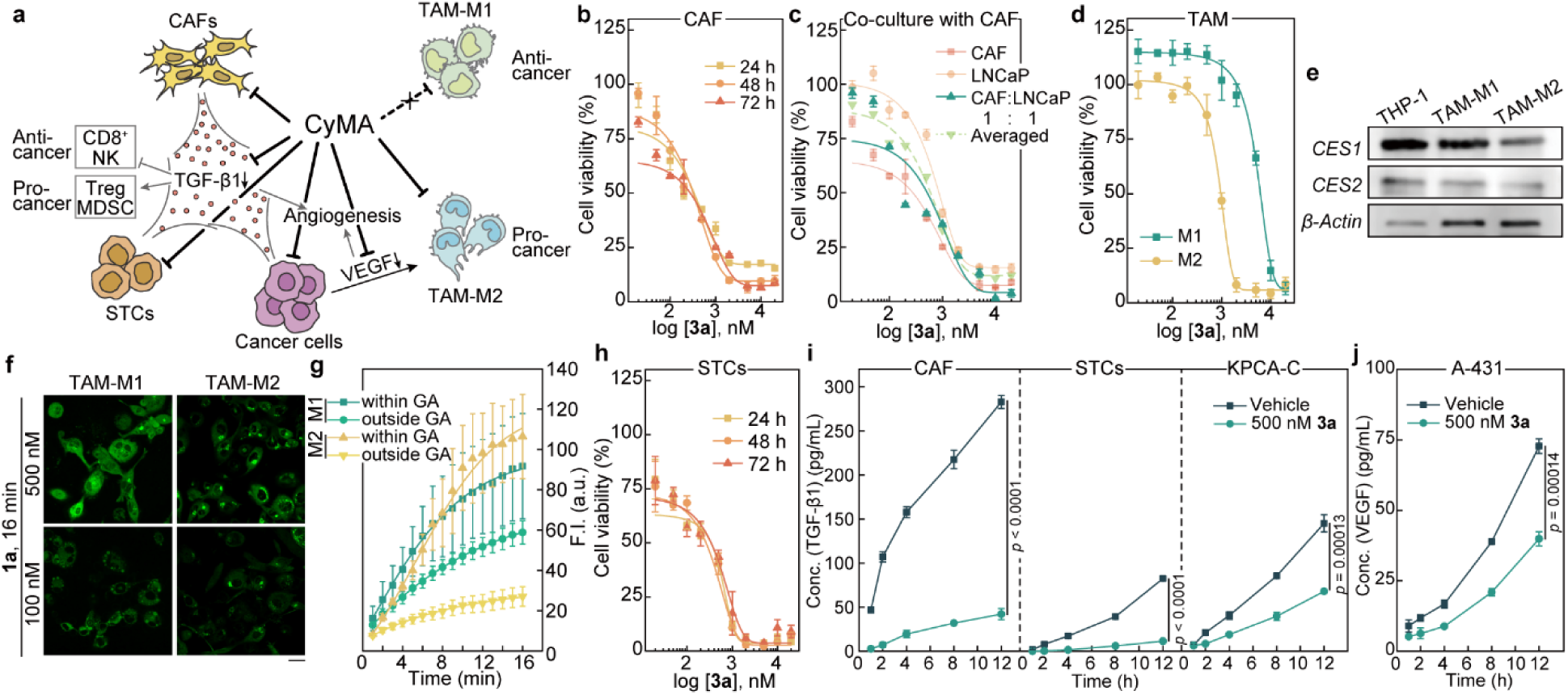
CyMA modulate tumor microenvironment. (a) Schematic illustration showing the inhibition of cancer associated fibroblasts (CAFs), senescent tumor cells (STCs), and selective inhibition of M2 tumor associated macrophages (TAMs) but not M1 TAM, as well as the suppression of cytokine secretion (TGF-β1 and VEGF). (b) Cell viability of CAFs treated with **3a** for 24, 48, 72 hours (n = 3). (c) Cell viability of the co-cultured CAFs with LNCaP cells treated with **3a** for 72 hours (n = 3). (d) Cell viability of M1 and M2 TAMs treated with **3a** for 24 hours (n = 3). (e) Immunoblotting of CES1, CES2 and β-Actin in THP-1 cells and THP-1 derived M1 and M2 TAMs. (f) CLSM images of M1 or M2 TAMs treated with **1a** (500 nM, 100 nM, 16 minutes). (g) Quantitative analysis of fluorescence intensity within or outside the Golgi in cells in (f) treated with **1a** (500 nM) (n = 8). (h) Cell viability of STCs treated with **3a** for 24, 48, 72 hours (n = 3). (i) Enzyme-linked immunosorbent assay (ELISA) of TGF-*β*1 in the conditioned media of CAFs, STCs, or KPCA-C cells after the treatment of vehicle or **3a** (500 nM) (n = 3). (j) ELISA of VEGF in the conditioned media of A-431 after the treatment of vehicle or **3a** (500 nM) (n = 3). Scale bar = 20 μm.

The CyMA precursor **3a** showed significant inhibitory effects on CAFs, with the IC_50_ decreasing to ∼300 nM after 72-h treatment (**Fig. 6b**). Using a co-culture model of patient-derived prostate cancer CAFs and LNCaP prostate cancer cells^89^, we observed that CyMA reduced cell viability more effectively than predicted by the average of individual cultures (**Fig. 6c**), suggesting that CyMA remodels the TME to limit tumor cell proliferation. In TAMs, CyMA displayed pronounced cytotoxicity against M2 macrophages^90^ (IC_50_ of 1 μM), while sparing M1 macrophages (IC_50_ > 6 μM) (**Fig. 6d**), presumably reflecting differences in CES expression, as confirmed by immunoblotting analysis (**Fig. 6e**). Notably, treatment with the fluorophore-conjugated CyMA precursor **1a** revealed distinct fluorescence patterns between macrophage subtypes. At 500 nM, **1a** accumulated at the Golgi of both M1 and M2 macrophages, but at 100 nM it selectively labeled the Golgi in M2 macrophages, whereas M1 macrophages presented dim spherical fluorescent structures (**Fig. 6f, g**). These results suggested that CES-mediated hydrolysis in M1 macrophage prevents Golgi accumulation, thereby reducing CyMA’s cytotoxicity.

Beyond CAFs and TAMs, CyMA potently inhibit chemotherapy-induced STCs (**Fig. 6h**; **Supplementary Fig. S11a**), which are known to promote metastasis via the senescence-associated secretory phenotype (SASP) and confer chemotherapy resistance^91^. Because CAFs and STCs support tumor growth through cytokine secretion, we quantified the TGF-β1 levels in conditioned media from CAFs, STCs, and cancer cells after CyMA (**3a**, 500 nM) or vehicle (DMSO) treatment. Within four hours, CyMA significantly reduced TGF-β1 secretion by all tested cell types (**Fig. 6i**). Importantly, secretion of FST—highly expressed by KPCA-C^92^— remained unaffected (**Supplementary Fig. S11b**), indicating that decreased TGF-β1 release was not a consequence of cell death. VEGF secretion was also lowered by CyMA treatment (**Fig. 6j**). Collectively, these findings highlight CyMA’s potential to broadly modulate the TME, laying the groundwork for combined strategies with ICB and other immunotherapies to achieve more effective cancer control.

## Summary

This study establishes CyMA as a first-of-its-kind supramolecular entity that actively and selectively target the Golgi, forming transient assemblies from enzymatically lipidated small molecules (< 700 Da) to trigger potent cellular responses with broad potential applications. By exploiting dynamic thioester bond formation and cleavage^93^, CyMA achieve cyclical and efficient activity at lower precursor concentrations than conventional assemblies formed by a single enzyme class. Although inhibitors against specific Golgi proteins are under development as anticancer agents^94,95^, they often face resistance due to protein mutations and adaptive feedback loops^96^. In contrast, CyMA formation relies on conserved biochemical reactions that generate non-diffusive aggregates at the Golgi, thereby disrupting intracellular trafficking and protein modification in a manner that affects multiple signaling pathways, including RTKs-AKT and RAS-ERK, and proving particularly effective against RAS-driven cancers. The heightened sensitivity observed in RAS-overexpressing FT33 and FT190 cells, as evidenced by their lower IC_50_ and IC_90_ values (**Fig. 3d,e**), underscores the broad-spectrum activity of CyMA and their reduced propensity for resistance. Although some RAS-driven cells exhibited ERK1/2 activation following CyMA treatment, it remains to be determined whether this contributes to survival or synthetic lethality. Importantly, CyMA also disrupt the cellular secretome by targeting Golgi trafficking, affecting both cytokine secretion and receptor localization. This dual mechanism— evident in decreased VEGF secretion (**Fig. 6j**) and VEGFR2 mislocalization (**Fig. 5g**)—prevents compensatory feedback and resistance often associated with selective cytokine blockade. Moreover, CyMA diminish immunosuppressive cytokines such as TGF-β, VEGF, and IL-10 (**Supplementary Fig. S11c**), potentially reshaping the tumor microenvironment, especially in contexts where RAS and TGF-β promote metastasis^97^. While extensive molecular engineering has linked CyMA molecular structures to Golgi-targeting properties (**Fig. 1c**, **Fig. 2o**), details of how CyMA disrupt Golgi-related trafficking remain to be elucidated. RNA-seq findings reveal differential Rab protein expression (e.g., Rab3d, Rab26, Rab27a/b, and Rab36 (**Supplementary Table S3**)) and are consistent with RUSH assays, suggesting that CyMA’s modulation of palmitoylation and vesicular trafficking could provide versatile tools for biological discovery and drug development.

## Methods

### Cell culture

T98G, U87MG, SH-SY5Y, MCF-7, H-460, U2OS, A-431, B16F10, VCaP, PC-3, HeLa, HEK293, NIH3T3, HepG2, RAW264.7, THP-1, SKOV-3, OVCAR-4, hTERT PF179T CAF cells were purchased from ATCC. KPCA-A, KPCA-B and KPCA-C cells were provided by Dr. Daniela Dinulescu lab. Saos-2 cells were provided by Prof. David Loeb lab. FT33, FT33+RAS, FT33+MYC, FT190, FT190+RAS, FT194, FT194+YAP, FT246, FT246+YAP, FT282, FT282+CCNE1 were provided by Dr. Ronny Drapkin lab. Cell lines were authenticated by CellCheck 9 - human (9 Marker STR Profile and Inter-species Contamination Test, IDEXX), confirming 100% match of the cell identity. T98G, MCF-7, HeLa, HEK293, HepG-2 cells were cultured in MEM supplemented with 10% FBS. SH-SY5Y cells were cultured in 1:1 mixture of EMEM and F12 Medium supplemented with 10% FBS. A-431, B16F10, VCaP, NIH3T3, RAW264.7 cells were cultured in DMEM supplemented with 10% FBS. H-460, Saos-2, THP-1, OVCAR-4 cells were cultured in RPMI 1640 medium supplemented with 10% FBS. U2OS, SKOV-3 cells were cultured in McCoy’s 5A medium supplemented with 10% FBS. U-87 MG cells were cultured in EMEM supplemented with 10% FBS. MCF-7 cells were cultured in EMEM supplemented with 10% FBS and 0.01 mg/mL human recombinant insulin. PC-3 cells were cultured in F-12K supplemented with 10% FBS. hTERT PF179T CAF cells were cultured in EMEM supplemented with 10% FBS and 1 µg/mL puromycin. KPCA-A, KPCA-B and KPCA-C cells were cultured in mFT cell media. FT33, FT33+RAS, FT33+MYC, FT190, FT190+RAS, FT194, FT194+YAP, FT246, FT246+YAP, FT282, FT282+CCNE1 cells were cultured in DMEM/F12 50:50 Mix without L-glutamine, supplemented with 10% FBS. All the cell lines were supplemented with 100 U/mL penicillin and 100 µg/mL streptomycin and were cultured and humidified with 5% CO_2_ at 37 °C.

### Cell transfection siRNA transfection

Cells were seeded at 2×10^5^ cells per well on the 6-well plate for 24 hours to allow attachment. Upon reaching 60-70% confluency, the cells were transfected with Lipofectamine 3000 reagent. Specifically, 5 µL of Lipofectamine 3000 and 5 µL of siRNA stock solution (20 µM) were separately dissolved in 250 µL of Opti-MEM and briefly vortexed before mixing. The mixture solution was left to stand for 15 minutes. Meanwhile, the media in each well was rinsed and replaced with 1.5 mL of Opti-MEM. The Lipofectamine 3000 and siRNA mixture was added dropwise to each well and incubated for 6 hours at 37 °C, after which the Opti-MEM was replaced with fresh culture media containing FBS and P/S. The cells were then incubated for 48 hours before proceeding with live-cell imaging or cell lysis for further analysis.

### Plasmid transfection

Cells were seeded at 1.5×10^5^ cells per confocal dish for 24 h to allow attachment. Upon reaching 40-50% confluency, the cells were transfected with Xfect™ Transfection Reagent. Specifically, 5 µg of the plasmid DNA was diluted with Xfect Reaction Buffer, followed by the addition of 1.5 μL Xfect Polymer. The mixture was briefly vortexed and incubated for 10 min at room temperature. The entire 100 μL of nanoparticle complex solution was then added dropwise to the cell culture medium, and the dish was rocked briefly. The confocal dish was incubated at 37°C overnight, and the media was replaced with fresh culture media for an additional 48-hour incubation. The cells were then ready for live-cell imaging.

### Confocal microscopy

A confocal dish (35 mm dish with 20 mm bottom well, #1.5 glass) was used to prepare CLSM samples. For live-cell imaging, cells in exponential growth phase were seeded on the confocal dish at 1.0×10^5^ cells per dish and incubated for 24h. After removing the culture medium, fresh medium containing the compound of interest was added to the cells for co-incubation at 37 °C in a humidified atmosphere of 5% CO_2_ for the desired period. Afterwards, the nuclei of cells were stained with Hoechst 33342 for 10 minutes, and the samples were washed with 1 mL of Live Cell Imaging Solution four times to fully remove the residual Hoechst 33342.

For time-lapse live-cell imaging, cells in exponential growth phase were seeded on a confocal dish at 1.0×10^5^ cells per dish and incubated for 24 h. The samples were washed with 1 mL of Live Cell Imaging Solution three times, and the nuclei were stained with Hoechst 33342 for 10 minutes. The samples were then washed with 1 mL of Live Cell Imaging Solution four times to remove the residual Hoechst 33342. The position of cells and the focal plane of the laser beam were determined using the fluorescence from the stained nuclei with a laser with 405 nm wavelength, and the Nikon Perfect Focus System was activated to prevent focus drift. The imaging solution in the confocal dish was replaced with fresh imaging solution containing the compound of interest. CLSM images of different channels were then recorded, with the time-series interval set to be 1 minute or no-delay. After a specified number of imaging cycles, the fluorescence images from both channels were saved for further analysis.

### Colocalization study with GALNT2-RFP

HeLa cells (1.5×10^5^ cells) were seeded in a confocal dish for 24 hours to allow attachment. The culture media was then replaced with fresh medium containing CellLight™ Golgi-RFP, BacMam 2.0 at the concentration of 2 µL per 10,000 cells, and the cells were incubated for another 24 hours to complete the transduction. The medium was then replaced with fresh media containing **1a** (1 µM) and incubated for 10 minutes for live-cell imaging. The person’s R value was calculated using Coloc 2 plugin in Fiji.

### Cell pretreated with inhibitors

HeLa cells (1.5×10^5^ cells) were seeded in a confocal dish for 24 h to allow attachment. The culture media was ten replaced with fresh medium containing endocytosis inhibitors (mβCD, EIPA, CPZ, Dynasore, 7-keto-chol), or inhibitors for LYPLA1/2 (ML211), PPT1 (DC661), palmitoylacyltransferases (2-BP), CES1 (Nevadensin), or CES2 inhibitor (Loperamide), and the cells were incubated for 30 minutes at 37 °C. Afterward, the cell medium was replaced with fresh medium containing CyMA for live-cell imaging.

### FRAP assay

FRAP was performed on a Zeiss LSM 880 confocal microscopy using a 63×/1.4 Oil objective or a Nikon AX-R resonant confocal system using a 60×/1.4 Oil objective. Five pre-bleach images were captured, followed by photobleaching using 488-nm laser at 100% intensity within a selected region. Another region was imaged without photobleaching as an internal control. 512×512-pixel images were captured at 0.26 s (or 5.02 s for Nikon AX-R CLSM) intervals using a 488-nm laser at 100% intensity with the pinhole set at 1 airy unit. Imaging continued until no further recovery was observed. The fluorescence recovery of the photobleached region was normalized and fitted into an exponential function.

### Structured illumination microscopy (SIM) imaging

A 3D-Nikon structured illumination microscopy (N-SIM, version AR5.11.00 64bit, Tokyo, Japan), equipped with solid-state lasers (488 nm, 561 nm, 640 nm, the output powers at the fiber end: 15 mW) and an Apochromat 100×/1.49 numerical aperture oil-immersion objective lens, was used to acquire all SIM images. Images were obtained using Nikon NIS-Elements 512 × 512 resolution, with Z-stacks. NIS-Elements AR Analysis was used to reconstruct and process raw images.

Cells were seeded on glass-bottomed culture dishes (MatTek; P35G-1.5-14-C) for 24 hours to allow for adhesion. For Golgi staining, cells were treated with **1a** (2 μM) for 5 min. Before imaging, cells were washed with PBS three times. Green channel images (emission bandwidth: 500-550 nm) were excited with a 488 nm laser, and red channel images (emission bandwidth: 570-640 nm) were excited with a 561 nm laser. Imaging data analysis was performed using ImageJ. Mitochondrial and ER analysis were conducted as per previously reported references^98,99^.

### ER-to-GA anterograde trafficking analysis

HeLa cells seeded on confocal dishes were transfected with various RUSH plasmids. The cells were then treated with CyMA for 6 hours at 37 °C, followed by nuclear staining with Hoechst 33342. The cells were transferred to the CLSM for imaging, and the medium was replaced with fresh medium containing 40 µM of biotin. Imaging began immediately after the addition of biotin using time-series mode. CLSM images were saved for further analysis.

### GA-to-PM anterograde trafficking analysis

HeLa cells seeded on the confocal dishes were transfected with various RUSH plasmids. Afterward, cells were treated with 40 µM of biotin for 1 hour at 20 °C to allow cargo proteins to accumulate at Golgi but inhibit their subsequent trafficking to the plasma membrane. Cells were then treated with CyMA for 6 hours at 20 °C. The cells were transferred to CLSM sites, and the temperature was raised to 37 °C. Imaging of the cells was performed immediately after increasing the temperature using time-series mode. The CLSM images were saved for further analysis.

### PM-to-GA retrograde trafficking analysis

HeLa cells seeded on the confocal dishes were treated with CyMA at different concentrations for 6 hours at 37 °C. Then, the media was replaced with fresh media containing 1 µg/mL CTxB-AF647 and incubated for another 1 hour. The cells were sent for CLSM imaging to study the localization and fluorescence intensity of CTxB-AF647 to evaluate the PM-to-Golgi trafficking behavior.

### GA-to-ER retrograde trafficking analysis

GA-to-ER trafficking was evaluated by photobleaching the ER pool of GalT and monitoring fluorescence recovery at the ER as an indicator of retrograde transport from Golgi to ER. HeLa cells transfected with GalT-EGFP were seeded in confocal dishes and incubated with CyMA for 6 hours at 37 °C. Afterward, FRAP of the ER pool of GalT was performed as described above. Time-series fluorescence intensity recoveries at the photobleaching sites were recorded for further analysis.

### Immunocytochemistry

The culture media was removed from cells, followed by two washes with PBS. Cells were fixed with 4% paraformaldehyde for 10 minutes and permeabilized with 0.1% Triton X-100 in PBS for 6 minutes. Cells were then blocked with 3% BSA and 22.5 mg/mL of glycine in PBST for 1 hour at room temperature. Primary antibodies were diluted 1:200 in 1% BSA in PBST and incubated with the cells overnight at 4 °C. Cells were then incubated with Alexa Fluor 647-conjugated secondary antibodies (1:1000 dilution) for 1 hour at room temperature. Between each step, except after blocking, cells were washed three times with PBS. The cells were then ready for CLSM imaging.

### Quantification of fluorescence intensity at Golgi

Image processing was conducted to extract single-cell responses. The image pixel values are scaled to be between [0, 1] with 1 indicates the original intensity of 255.

a. Nuclei Detection. The Otsu method^100^ was used on nucleus-staining frames to automatically determine an intensity threshold for detecting foreground pixels corresponding to nuclei. The minimum threshold was set to 0.6 out of 1.0. Morphological operations (open, fill, and erode) were applied to remove noise, fill the holes in the foreground segments, and separate foreground pixels into segments corresponding to individual nuclei. Some nuclei near image boundaries were discarded.
b. Reaction Signal Detection. The last reaction-staining frame of a video was used to determine the intensity threshold for detecting reaction signals in the entire video. The threshold was set to a value higher than 99% of the pixels in the last frame. Small foreground segments containing less than 4 pixels were considered noise and removed. A foreground segment was considered part of a cell if its corresponding nucleus were the closed to it.
c. Golgi Detection. A manual threshold of 0.05 was set to determine the foreground mask representing Golgi signals. Using the Golgi mask, reaction signals were classified as occurring inside or outside the Golgi.

### Palmitoylation of CyMA characterized by LC/HR-MS

A total of 1.2×10^7^ cells were treated with 1 µM of CyMA or vehicle (DMSO) for 30 minutes, and then washed with HEPES buffer twice. Cells were collected and centrifuged to obtain a pellet. The pellet was resuspended in 500 µL of HEPES buffer, followed by the addition of 1.5 mL of DCM and 1 mL of methanol. The mixture was allowed to sit for 10 minutes. Afterward, 0.5 mL DCM and 0.5 mL Tris-HCl (50 mM, pH 2.0) were added, and the tube was centrifuged to collect the organic phase. The organic phase was washed with buffer (1 mL methanol+1 mL Tris-HCl (50 mM, pH 2.0)) and centrifuged again to obtain the organic phase. The organic solvent was evaporated with N2, and the remaining solid was dissolved in 150 µL of methanol and sent for LC/HR-MS analysis.

### MTT assay

The MTT assay was used to determine cell viability for cytotoxicity evaluation. Cells were seeded at 1×10^4^ cells per well in 96-well plates for 24 h to allow attachment. Culture media were replaced with fresh culture media containing the compounds at a series of concentrations. After 24, 48, and 72 hours, 10 µL of MTT solution (5 mg/mL) was added to each well, and the plate was incubated in the dark for 4 h at 37 °C. Then, 100 µL of 10% SDS-HCl was added to stop the reaction and dissolve the formazan. The absorbance at 595 nm was determined by a microplate reader. The assay was repeated three times, and the mean values of three measurements were plotted, with error bars representing standard deviation. For 7-day cytotoxicity, seed cells at 5,000 cells per well, and fresh D-peptide-containing medium was added every 3 days.

### Immunoblotting

Cells were cultured to 80-90% confluency, lysed with 500 µL lysis buffer (containing protease inhibitor cocktail and phosphatase inhibitor cocktails) per 10 cm dish on ice, sonicated for 10 seconds, and subjected to three freeze-thawed cycles to collect cell lysates from various cell lines.

The lysates were centrifuged at 12,000 rpm for 10 minutes at 4 °C, and the supernatant was collected. The proteins in the lysates were denatured by adding sample loading buffer and incubating at 95 °C for 8 minutes. The lysate samples were loaded onto precast gels for electrophoresis at 140 V for 40 minutes, followed by blotting onto PVDF membrane under 100 V for 100 minutes in ice bath. Membranes were blocked with blocking buffer for 1 hour at room temperature and incubated with primary antibody (1:1000 dilution) at 4 °C overnight. After washing with TBST three times (5 min per wash), the membrane was incubated with secondary antibody (1:10,000 dilution) for 1 hour at room temperature. After washing with TBST six times, chemiluminescent substrate was added and incubated for 1 minute. The membranes were then scanned using a blot scanner. Subsequently, the scanned membranes were stripped using Restore™ Western Blot Stripping Buffer, blocked, and incubated with the next primary antibody.

### Cell migration assay

Cell migration performance was evaluated using Radius™ Cell Migration Assays, which provide a convenient method to measure two-dimensional cell migration. When cells are seeded in the well of the Radius plate, they adhere everywhere except in the center of the well, where a biocompatible hydrogel spot is placed. Once cells form a monolayer, the assay is initiated by gently dissolving the gel with a removal solution, leaving a gap across which cell migration can occur. Specifically, 1×10^5^ cells per well were seeded in Radius™ Migration Plate for 24 h to allow attachment. The media were then aspirated, and the wells were washed three times with 0.5 mL of fresh media. Afterward, the media were aspirated again, and 0.5 mL of 1× Radius™ Gel Removal Solution was added. The plate was transferred to a cell culture incubator for 30 minutes to allow complete gel removal. The wells were then washed three times with 0.5 mL of fresh media. After the final wash, 1 mL of complete medium containing CyMA or vehicle (DMSO) was added to each well. Pre-migration images were captured using an inverted microscope, and the images of migrating cells in each well were captured at different time points for further analysis.

### Cell spheroids generation

Spheroids were generated using a low-adhesion U-bottom microplate, specifically the Thermo Scientific™ Nunclon™ Sphera™ 96U-well microplate (Cat. No. 174929). KPCA-C cells (10,000 cells per 100 µL) were seeded into each well and incubated in a cell incubator for 48 hours to generate spheroids.

### 3D Cell Viability Assay

The cytotoxicity of CyMA against the generated cell spheroids was tested using The CellTiter-Glo® 3D Cell Viability Assay, which measures ATP as an indicator of viability and generates a luminescent readout that is much more sensitive than colorimetric or fluorescence-based methods. Specifically, 100 µL of CellTiter-Glo® 3D Reagent was added into each well containing 100 µL of culture media with cell spheroids. The contents were mixed vigorously for 5 minutes to induce cell lysis, followed by a 25-minute incubation at room temperature to stabilize the luminescent signal. Luminescence was recorded using a microplate reader.

### Patient-derived organotypic tumor spheroids (PDOTS)

Patient tissue studies were reviewed and approved by the Institutional Review Board of Brigham and Women’s Hospital. Patient-derived tumor ascites were collected by paracentesis and centrifuged to form a cell pellet. The supernatant was aspirated, and the pellet was resuspended in ACK lysis buffer (Thermo Fisher, A1049201). After maximum hemolysis was observed, PBS was added, and the process was repeated to ensure red blood cells and hemolytic products were removed. The final cell pellet was gently agitated to promote the resuspension into spheroids. The resuspended sample was filtered through a 100 μm filter. The supernatant was labeled as S2+3 while the filtered cell aggregates were labeled as S1, following previously established naming conventions^101^. On ice, a mixture of collagen, NaOH, phenol red, water, and PBS was made and adjusted to a pH of approximately 7.3-7.4. The spheroids were pelleted again by centrifuging at 300 g for 3 minutes. This pellet was resuspended in the collagen mixture, and 10 µL of the spheroid-collagen mixture was loaded into a microfluidic device (AIM Biotech, DAX- 1) as previously described^101,102^. The devices were incubated at 37°C for 35-40 minutes to allow the collagen to polymerize and form a matrix. After polymerization, 300 µL of RPMI media supplemented with 6,000 U/mL IL-2 (Miltenyi Biotec, 130097746) was divided evenly between all four ports with treatments added: 3 µM **3a**, 6 µM **3a**, 200 μg/mL anti-PD-1 (Fisher Scientific, 501360845) and anti-PD-L1 (Fisher Scientific, 501360846), or 3 µM of **3a** with 200 μg/mL anti-PD-1 and anti-PD-L1.

After 5-7 days, live-dead analysis was performed to determine cell viability in each treatment condition. A 1:1 dilution of AO/PI stain (Nexcelcom, CS2-0106) in PBS was made and 20-30 μL of this solution was added to each device. After 5 minutes, the devices were imaged using an inverted Nikon Eclipse Ti microscope equipped with Nikon DS-Qi1Mc camera using NIS-Elements software. Using this software, live (acridine orange (AO)) and dead (propidium iodide (PI)) cell areas were quantified for analysis.

### Drug resistance test

KPCA-B or KPCA-C cells were divided into two culture dishes. Group 1: The KPCA cells were incubated with **3a** (500 nM) for 24 hours, and the media was replaced with fresh media for 2 more days to allow the cells to expand. The cells were then subcultured and allowed to proliferate until they reached 80-90% confluency, followed by incubation with **3a** to initiate the next cycle of cell stimulation. Group 2: KPCA cells incubated with vehicle (DMSO) instead of **3a**, following the same procedure as a control. We seeded stimulated cells in from Group 1 and unstimulated KPCA cells (Group 3) in 96-well plates at 1×10^5^ cells/well for 24 hours. The cells were then treated with **3a** for 24 hours, and the cell viability was measured using the MTT assay.

### In vivo experiments

Animal studies were conducted following the guidelines provided by the Institutional Animal Care and Use Committee (IACUC) of Brigham and Women’s Hospital. Tumor engraftment was performed through intraperitoneal injection of 3.7 x 10^6^ KPCA-B cells resuspended in a 1:1 solution of PBS and Matrigel (Corning, 354234) into 6-week C57BL/6J female mice (Jackson Laboratories, Strain # 000664). Mice were treated via intraperitoneal injection with **3a** resuspended in PBS for a final circulating blood concentration of 3 µM (0.144 mg/kg), 50 µg of Anti-PD-L1 (BioCell, BE0101) resuspended in PBS at pH 6.5 (BioCell, IP0065), 50 µg of Anti-CTLA-4 (BioCell, BE0131) resuspended in PBS at pH 7.0 (BioCell, IP0070), or a combination of **3a** (0.144 mg/kg), 50 µg of Anti-PD-L1, and 50 µg of Anti-CTLA-4. Tumor burden was determined by collecting tumors through necropsy. Blood samples were obtained via cardiac puncture in EDTA coated microcentrifuge tubes (Owens & Minor, 0723365974), which were immediately centrifuged at 2000×g for 15 minutes at 4 °C. The plasma supernatant was collected and stored at -80 °C.

### Metabolic labeling

HeLa cells (1.5×10^5^ cells) were seeded in a confocal dish for 24 h to allow attachment. The next day, the media was replaced with fresh aliquots containing 50 µM of azido-metabolic molecule (e.g., palmitic acid, myristic acid, Ac4GlcNAc, Ac4GalNAc, Ac4ManNAc) in culture medium and add into the experimental and positive control cells. The cells were incubated for an additional 12 hours. After this, the media was replaced with fresh media containing either (CyMA + azid-metabolic molecule) or (DMSO + azido-metabolic molecule), and the cells were incubated for another 12 hours. The cells were then rinsed, fixed, and fluorophore (AF488- alkyne) was linked to the azido incorporated proteins using a standard click reaction. After washing the cells three times with PBS, they were imaged using CLSM to quantify fluorescence intensity.

### Immunoprecipitation of ubiquitinated proteins

KPCA-C cells were treated with CyMA and lysed. Ubiquitinated proteins were pulled down using Signal-Seeker ^TM^ Ubiquitination Detection Kit, following the manufacturer’s protocol. Both the input lysate and enriched ubiquitinated proteins were analyzed by immunoblotting.

### Proteomics

#### Proteomics of palmitoylated proteins

KPCA-C cells were metabolically labeled with azido-palmitic acid followed by CyMA treatment. Palmitoylated proteins were pulled down using Click-&-Go® Protein Enrichment Kit to capture azide-modified proteins according to the manufacturer’s protocol. On-bead digestions were stored in 50 mM Tris buffer and sent for LC-MS/MS analysis.

#### Proteomics of ubiquitinated proteins

Enriched ubiquitinated proteins from KPCA-C were loaded on precast gels and run under electrophoresis at 100 V for 10 minutes. The corresponding gel sections were cut out and sent for LC-MS/MS analysis.

#### Proteomics of pulled down proteins from UV-crosslinking

1.2×10^7^ of HeLa cells were treated with or without **1k** (5 μM, 12 mL) for 30 minutes. After removing the media containing **1k**, cells were washed with PBS buffer three times. Cells with or without the pretreatment were placed in the chamber of Stratagene Stratalinker 2400 UV Crosslinker, and underwent a UV-crosslinking (365 nm, 15 watts × 6, 1 min). The cells were then washed once with PBS buffer and digested with trypsin, followed by centrifugation (1500 rpm, 3 minutes). The cell pellets were washed with PBS buffer three times (1500 rpm, 3 minutes). The cells were then suspended in 1 mL of TBST with 1% protease inhibitor cocktail and lysed by four freeze-thaw cycles. 2 μL of NBD antibody was added to the cell lysate and co-incubated overnight at 4 °C. Afterwards, 2 μL of goat anti-rabbit IgG (H+L) secondary antibody and biotin were added and incubated at room temperature for 1.5 hours. 50 µL (0.5 mg) of Pierce Streptavidin Magnetic Beads were added to a 1.5 mL microcentrifuge tube and placed on a magnetic stand to collect the beads. The beads were washed with TBST (1 mL) three times. The cell lysate solution was added to the pre-washed beads and incubated for 1.5 h at room temperature with mixing. The beads were then washed with TBST (1 mL) for three times, and 100 µL of Elution Buffer (IgG Elution Buffer, pH 2.0) was added. The tube was incubated at room temperature with mixing for 5 minutes, after which the beads were magnetically separated. The supernatant containing POI was saved, lyophilized, and sent for LC-MS/MS analysis.

#### Induction of M1 and M2 macrophage

M1 and M2 macrophage cells were differentiated from THP-1 monocytes as previously reported^103^. Specifically, human THP-1 monocytes were differentiated into macrophages by incubating them in the presence of 150 nM of phorbol 12-myristate 13-acetate (PMA). The resulting M0 macrophage cells were polarized into M1 macrophages by incubation with IFN-γ in combination with LPS. M2 macrophages were induced in vitro by IL-4 and IL-13 stimulation.

#### Induction of senescent cells

HeLa cells were grown to 80% confluency and treated with 1 μg/mL of cisplatin for 24 hours. After treatment, the cells were rinsed twice with PBS and cultured for an additional 6-7 days in fresh complete media, with media change every 2-3 days. The senescent HeLa cells exhibited abnormal morphology and increased SA-βGal expression level.

#### ELISA

Cytokine secretion in the conditioned media was assayed using an ELISA kit, following the procedure recommended by the supplier. Opti-MEM was used as the media to avoid the external source of TGFB1 or other cytokines from FBS. TGFB1 proteins were activated according to the manufacturer’s protocol before being applied to ELISA.

#### Quantification and Data Analysis

All graphs were created using Origin 2021. Quantification of fluorescence intensity was conducted using Fiji. Error bars represent s.d. unless otherwise noted. For comparison between two groups, *p* values were determined using two-tailed Student’s t-tests. All cell-based or cell-free experiments were repeated two to three times.

#### Data and materials availability

All data are available in the main text or the supplementary materials.

## Acknowledgements

We thank Havard Taplin Mass Spectrometry Facility for conducting LC-MS/MS analysis. This work is partially supported by NIH CA142746 (BX), EY036512 (BX), NSF DMR-2011846 (BX), Ministry of Education (Singapore) Tier 2 MOE-T2EP30221-0001 (LL).

## Author contributions

W.T. and B.X. designed the project, synthesized the compounds, cultured cells, analyzed the data, and wrote the manuscript, with input from all authors. Q.Z. and Z.L. synthesized the compounds and cultured cells. K.Q. and J.D. assisted the SIM experiments. D.M. and L.L. assisted cell experiments. T.G., D.D. and J.H. assisted the *ex vivo* and *in vivo* experiments. N.C., D.L. and R.D. assisted the cell culture and analysis of cytotoxicity. C.X. and C.L. assisted the proteomic analysis. W.L., M.L. and I.A. assisted the cell experiments. P.H. assisted the image processing.

## Supplementary Materials

Materials and Methods

Supplementary Figures: Supplementary Figs. S1-S19

Supplementary Tables S1-S3 References

Supplementary Movie S1

