## Supplementary Materials for "Cycling Molecular Assemblies for Selective Cancer Cell Golgi Disruption"

**Materials and Methods**

**Materials**

2-Cl-trityl chloride resin (1.0 mmol/g), Fmoc protected amino acid, and HBTU were obtained from GL Biochem (Shanghai, China). N, N-diisopropylethylamine (DIEA), 2-Mercaptoethanol, chloroquine diphosphate salt, and solvents were obtained from Fisher Scientific. 4-Chloro-7-nitrobenzofurazan was obtained from Alfa Aesar. β-alanine was purchased from Indofine Chemical Company. 1-[(2-Aminoethyl)sulfanyl]ethan-1-one hydrobromide was purchased from Sigma-Aldrich. Ethylene glycol, acetyl chloride, 2-naphthoic acid, acetoxyacetyl chloride were purchased from TCI America. Biphenyl-3-carboxylic acid, [1,1'-biphenyl]-2-carboxylic acid, 4-biphenylacetic acid, 4-(4-pyridyl)benzoic acid, 4-pyridin-3-yl-benzoic acid, 4-(2-pyridyl)benzoic acid, 2,2'-bipyridine-5-carboxylic acid were purchased from 1PlusChem. 4-Benzylbenzoic acid was purchased from Enamine. Methyltrioxorhenium(VII) was purchased from Sigma-Aldrich. Fmoc-D-2,3-diaminopropionic acid was purchased from AKSci. 4'-Hydroxy-[1,1'-biphenyl]-4-carboxylic acid, methyl 3-chloro-3-oxopropanoate were purchased from AmBeed. Nevadensin, loperamide hydrochloride and DC661 were purchased from MedChemExpress. Cisplatin, dynasore, 3-methyladenine, wortmannin, thapsigargin, temsirolimus were purchased from Selleckchem. ML-211 was purchased from APExBIO. 7-Keto cholesterol was purchased from Cayman. Lipopolysaccharides, phorbol 12-myristate 13-acetate, tunicamycin and 2-bromohexadecanoic acid were purchased from Sigma-Aldrich. All the chemical reagents and solvents were used as received from commercial sources without further purification.

Nitrobenzofurazan polyclonal antibody (Catalog # PA1-85035), goat anti-rabbit IgG (H+L) secondary antibody, biotin (Catalog # 65-6140) were purchased from Invitrogen. Pierce™ Streptavidin Magnetic Beads (Catalog # 88816), Pierce™ IgG Elution Buffer, pH 2.0 (Catalog # 21028) were purchased from Thermo Scientific. Human PPT1 siRNA (Catalog # abx904208), Human LYPLA1 siRNA (Catalog # abx903086), Human LYPLA2 siRNA (Catalog # abx903087) were purchased from abbexa. Human ZDHHC3 siRNA (Catalog # AM16708), Human ZDHHC7 siRNA (Catalog # AM16708), Human ZDHHC12 siRNA (Catalog # AM16708), Human ZDHHC20 siRNA (Catalog # AM16708) were purchased from ThermoFisher Scientific. Lipofectamine 3000 Transfection Reagent (Catalog # L3000008), Silencer™ Negative Control No. 1 siRNA (Catalog # AM4611) were purchased from Invitrogen. Xfect™ Transfection Reagent (Catalog # 631317) was purchased from TaKaRa. Recombinant human IFN-gamma protein (Catalog # 285-IF), recombinant human IL-4 protein (Catalog # 204-IL), recombinant human IL-13 protein (Catalog # 213-ILB) were purchased from Bio-Techne Corporation. Cholera toxin subunit B (Recombinant), Alexa Fluor™ 647 conjugate (Catalog # C34778) was purchased from Invitrogen. TWEEN® 20, and Immobilon™-FL PVDF membranes (catalog # IPFL00005) were purchased from Millipore Sigma.

CellLight™ Golgi-RFP, BacMam 2.0 (Catalog # C10593) and CellLight™ Lysosomes-RFP, BacMam 2.0 (Catalog # C10597) were purchased from Invitrogen. LYPLA1 polyclonal antibody (Catalog # PA5-101405), LYPLA2 polyclonal antibody (Catalog # PA5-27653), PPT1 polyclonal antibody (Catalog # PA5-29177), CES1 polyclonal antibody (Catalog # PA5-19740), CES2 polyclonal antibody (Catalog # PA5-102415), N-Ras Recombinant rabbit monoclonal antibody (Catalog # 703435), mTOR polyclonal antibody (Catalog # PA5-34663), TGN46 polyclonal antibody (Catalog # PA5-23068), GAPDH polyclonal antibody (Catalog # PA1-16777), CXCR4 recombinant rabbit monoclonal antibody (Catalog # 704015), VEGF receptor 2 monoclonal antibody (Catalog # MA5-15157), Phospho-ERK1/ERK2 polyclonal antibody (Catalog # 44-680G), ERK1/ERK2 monoclonal antibody (Catalog # MA5-15134), Phospho-p38 MAPK monoclonal antibody (Catalog # MA5-15182), p38 MAPK recombinant rabbit monoclonal antibody (Catalog # MA5-41213), Phospho-JNK1/JNK2 polyclonal antibody (Catalog # 44-682G), EGFR polyclonal antibody (catalog # PA1-1110), goat anti-rabbit IgG (H+L) secondary antibody, HRP (catalog # 31460), goat anti-rabbit IgG (Heavy chain), superclonal™ recombinant secondary antibody, Alexa Fluor™ 647 (catalog # A27040), SuperSignal™west pico PLUS chemiluminescent substrate (catalog # 34579), SuperBlock™ (TBS) blocking buffer (catalog # 37535), Restore™ PLUS western blot stripping buffer (catalog # 46428), and Halt™ protease inhibitor cocktail (100X) (catalog # 78430) were purchased from Invitrogen. KRAS polyclonal antibody (Catalog # 12063-1-AP), HRAS polyclonal antibody (Catalog # 18295-1-AP), GNAQ polyclonal antibody (Catalog # 13927-1-AP), JNK polyclonal antibody (Catalog # 10023-1-AP), FGFR2 polyclonal antibody (Catalog # 13042-1-AP) were purchased from Proteintech. Goat anti-rabbit IgG H&L (Alexa Fluor® 647) (Catalog # ab150079), rabbit recombinant monoclonal LC3B antibody (Catalog # ab192890), anti-pan-AKT antibody (Catalog # ab8805), anti-pAKT antibody (Catalog # ab192623), anti-beta Actin antibody (Catalog # ab8227) were purchased from abcam. Insulin receptor β (4B8) rabbit mAb (Catalog # 3025S) was purchased from Cell Signaling Technology. Human TGF-beta 1 ELISA Kit (Catalog #: ELH-TGFb1-1), Mouse TGF-beta 1 ELISA Kit (Catalog # ELM-TGFb1-1), Mouse follistatin ELISA Kit (Catalog # ELM-FST-1), Human TGF-alpha ELISA Kit (Catalog # ELH-TGFa-1) were purchased from RayBiotech. Human VEGF ELISA Kit (Catalog # KHG0111) was purchased from Invitrogen. Sample Activation Kit 1 (Catalog # DY010) was purchased from Bio-Techne Corporation. Click-&-Go® Cell Reaction Buffer Kit (Catalog # CCT-1263), N-azidoacetylmannosamine-tetraacylated (Ac4ManNAz) (Catalog # CCT-1084), N-azidoacetylglucosamine-tetraacylated (Ac4GlcNAz) (Catalog # CCT-1085), N-azidoacetylgalactosamine-tetraacylated (Ac4GalNAz) (Catalog # CCT-1086), azido myristic acid (Catalog # CCT-1345), azido palmitic acid (Catalog # CCT-1346) were purchased from vector laboratories. AF 488 alkyne (Catalog # B18B0) was purchased from Lumiprobe.

ptfLC3 was a gift from Tamotsu Yoshimori (Addgene plasmid # 21074; http://n2t.net/addgene:21074; RRID:Addgene_21074). pEGFR-miRFP670nano3 was a gift from Vladislav Verkhusha (Addgene plasmid # 184673; http://n2t.net/addgene:184673; RRID:Addgene_184673). pCW57-CMV-ssRFP-GFP-KDEL was a gift from Noboru Mizushima (Addgene plasmid # 128257; http://n2t.net/addgene:128257; RRID:Addgene_128257). pCLBW cox8 EGFP mCherry was a gift from David Chan (Addgene plasmid # 78520; http://n2t.net/addgene:78520; RRID:Addgene_78520). ECFP-ELP1-25 was a gift from Michael Davidson (Addgene plasmid # 55341; http://n2t.net/addgene:55341; RRID:Addgene_55341). pmScarlet3-Giantin_C1 was a gift from Dorus Gadella (Addgene plasmid # 189773; http://n2t.net/addgene:189773; RRID:Addgene_189773). piRFP670-N1-GalT was a gift from Lei Lu (Addgene plasmid # 87325; http://n2t.net/addgene:87325; RRID:Addgene_87325). Str-Ii_SBP-EGFP-Golgin84 was a gift from Franck Perez (Addgene plasmid # 65303; http://n2t.net/addgene:65303; RRID:Addgene_65303). pEF.myc.ER-E2-Crimson was a gift from Benjamin Glick (Addgene plasmid # 38770; http://n2t.net/addgene:38770; RRID:Addgene_38770). Str-KDEL_ManII-SBP-mCherry was a gift from Franck Perez (Addgene plasmid # 65253; http://n2t.net/addgene:65253; RRID:Addgene_65253). pcDNA3-Golgi-ExRai-AktAR2 was a gift from Jin Zhang (Addgene plasmid # 184051; http://n2t.net/addgene:184051; RRID:Addgene_184051). Str-KDEL_TNF-SBP-mCherry was a gift from Franck Perez (Addgene plasmid # 65279; http://n2t.net/addgene:65279; RRID:Addgene_65279). Str-KDEL_SBP-mCherry-Ecadherin was a gift from Franck Perez (Addgene plasmid # 65287; http://n2t.net/addgene:65287; RRID:Addgene_65287).

Minimum Essential Medium (MEM), Dulbecco's Modified Eagle Medium (DMEM), McCoy's 5A, fetal bovine serum (FBS) Opti-MEM, CO_2_ Independent Medium and penicillin-streptomycin (PS) were purchased from Gibco. RPMI 1640 medium, F-12K medium, and Eagle's Minimum Essential Medium (EMEM) were purchased from American Type Culture Collection (ATCC, USA). Insulin from bovine pancreas was purchased from Sigma-Aldrich.

Radius™ 24-Well Cell Migration Assay was purchased from CELL BIOLABS, INC. Palmitoylated Protein Assay Kit (Green) (Catalog # ab273282) was purchased from abcam. Pierce™ Glycoprotein Staining Kit (Catalog # 24562), Mammalian beta-Galactosidase Assay Kit (Catalog # 75707), Pierce™ Silver Stain Kit (Catalog # 24612), Nunclon™ Sphera™ 96-Well, Nunclon Sphera-Treated, U-Shaped-Bottom Microplate (Catalog # 174929) were purchased from Thermo Scientific. Signal-Seeker™ Ubiquitination Detection Kit (Catalog # BK161) was purchased from Cytoskeleton, Inc. CellTiter-Glo® 3D Cell Viability Assay (Catalog # G9681) was purchased from Promega.

**Instruments**

All precursors and compounds were purified by a reverse phase HPLC (Agilent 1100 Series) equipped with an XTerra C18 RP column. HPLC grade acetonitrile (0.1% TFA) and HPLC grade water (0.1% TFA) were used as the eluents. LC-MS spectra were obtained with a Waters Acquity Ultra Performance LC with Waters MICROMASS detector and a Bruker Elute PLUS UHPLC with a Bruker timsTOF Pro. Fluorescence images were captured using a ZEISS LSM 880 confocal laser scanning microscope, a ZEISS LSM 880 AiryScan Fast Confocal System, and a Nikon AX-R Resonant Confocal System.

**Methods**

**Synthesis of NBD-β-Alanine**

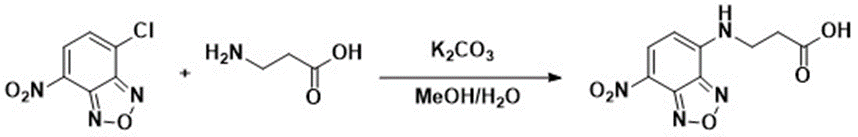

**Scheme S1.** Synthetic procedure of NBD-β-Alanine.

To a 10 mL aqueous solution of β-Alanine (5.5 mmol, 490 mg) and potassium carbonate (16.5 mmol, 2.07g), NBD-Cl (5 mmol, 1g) in 60 mL of MeOH was added dropwise with stirring under nitrogen gas protection. After stirring at room temperature for 6 hours, the methanol was removed by a rotary evaporator and the residual solution was acidified to pH 3 using 1N HCl. The acidic aqueous solution was then extracted by diethyl ether. The combined organic solution was dried over anhydrous sodium sulfate and concentrated by rotary evaporator. The resulting dark-yellow powder (NBD-β-Alanine) was used directly for solid phase peptide synthesis.

**CyMA synthesis**

**Synthesis of 2a, 5a**

We used standard Fmoc chemistry for solid phase peptide synthesis (SPPS) with 2-chlorotrityl chloride resin and Fmoc-protected amino acids with appropriately protected side chains. Briefly, the 2-Cl resin (1 g) was swelling in dry DCM for 30 minutes, followed by loading the first amino acid onto the resin. Next, 10 equivalents of 2-mercaptoethanol dissolved in DCM/DMF (v:v=1:1) were added into the SPPS reactor and incubated with the resin overnight at room temperature. The thiol group of 2-mercaptoethanol is selectively attached to the resin due to its high nucleophilicity. After discarding the liquid phase, the resin with mercaptoethanol linker was washed five times with DMF. Subsequently, Fmoc-protected amino acid (1.5 equiv.), along with N,N′-dicyclohexylcarbodiimide (DCC, 2 equiv.) and a catalytic amount of 4-Dimethylaminopyridine (DMAP) dissolved in DMF, was added to the reactor and incubated with the resin overnight at room temperature. The following day, the Fmoc group was removed with 20% piperidine in DMF, and the next Fmoc-protected amino acid was coupled to the free amino group using HBTU as the coupling reagent. The N-terminus of the peptide was capped with different NTGs on the resin to give various CyMA. The peptide chain was cleaved from the resin by 95% TFA (95% TFA, 2.5% TIPS, 2.5% H_2_O) for 1 hour. After solvent removal via rotary evaporation, 30 mL of dry diethyl ether was added to the residual solution, followed by centrifugation at 5000 rpm for 8 minutes. The resulting solid products were dried by lyophilization and further purified with RP-HPLC (**Scheme S2**). Capital letter represents L-configurational amino acids, while lowercase letter represents D-configurational amino acids (e.g., F = L-phenylalanine; f = D-phenylalanine).

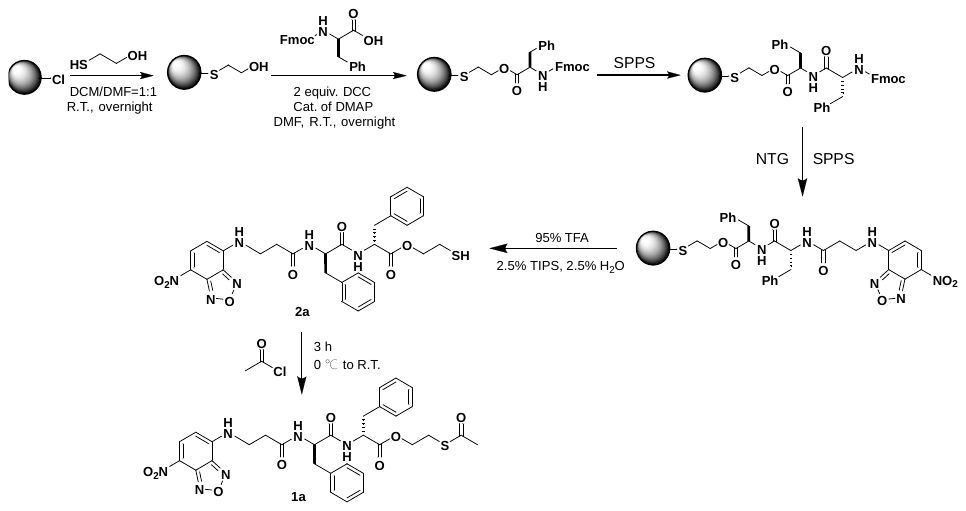

**Scheme S2.** Synthetic procedure of the representative CyMA, **2a** and CyMA precursor, **1a**.

**CyMA precursors synthesis**

The thiol group on 2-mercaptoethanol is selectively attached to the resin due to its high nucleophilicity. An ester bond is formed between the hydroxyl group from 2-mercaptoethanol (linked on the resin) and the carboxylic group of the Fmoc-protected amino acid by Steglich esterification. After standard solid-state peptide synthesis (SPPS), the derived peptide thiol, cleaved from the resin, is directly reacted with an acyl chloride in TFA to generate a thioester in high yield. The crude CyMA precursors are further purified by reverse-phase HPLC to yield the final CyMA precursors.

**Synthesis of 1a-1v, 3a-3r, 4a-4d**

The synthesized CyMA were dissolved in dry TFA and chilled in iced water bath and flushed with nitrogen. Then, 4 equivalents of the according acyl chloride were added dropwise into the TFA solution with CyMA, and the temperature was raised to room temperature. The reaction was left standing for 3 hours and then quenched with ice-cold water. After the solvent was removed by rotary evaporation, the resulting oily product was purified directly by RP-HPLC to give the CyMA precursors with thioester bonds (**Scheme S2**).

**Synthesis of 6, 7 as analogues of 1a, 3a with amide bonds**

Briefly, 0.1 mmol of the synthesized peptide (NBDff for **6**, BPff for **7**) was dissolved in 2 mL dry DMF, and 0.12 mmol of HBTU was added directly to the solution. After stirring the mixture for 30 minutes, 0.12 mmol of 1-[(2-aminoethyl)sulfanyl]ethan-1-one hydrobromide was added, and DIEA was added dropwise to adjust pH around 8. The reaction mixture was stirred for 8 hours. Afterward, the solvent is air dried, and the remained oily product was dissolved in methanol and purified by RP-HPLC to obtain the corresponding titled compounds.

**Synthesis of controls**

**Synthesis of 2b, 2c, 5b, 5c**

2-Cl resin (1 g) was swelling in dry DCM for 30 minutes, followed by loading the first amino acid onto the resin. Next, 10 equivalent of ethylene glycol dissolved in DCM/DMF (v:v=1:1) mixed with 3 equivalents of DIEA was added to the SPPS reactor and incubated with the resin overnight at room temperature. The liquid phase was discarded, and the resin with ethylene glycol linker was washed five times with DMF. Subsequently, Fmoc-protected amino acid (1.5 equiv.), N,N′-dicyclohexylcarbodiimide (DCC, 2 equiv.), and a catalytic amount of 4-Dimethylaminopyridine (DMAP) dissolved in DMF was addedto the SPPS reactor and incubated with the resin overnight at room temperature. The following day, Fmoc group was then removed with 20% piperidine in DMF, and the next Fmoc-protected amino acid was coupled to the free amino group using HBTU as the coupling reagent. The N-terminus of the peptide was capped with different NTGs on the resin to give CyMA. The peptide chain was cleaved from the resin by 95% TFA (95% TFA, 2.5% TIPS, 2.5% H2O) for 1 hour. After solvent removal via rotary evaporation, 30 mL of dry diethyl ether was added to the residual solution, followed by centrifugation at 5000 rpm for 8 minutes. The resulting solid products were dried by lyophilization and further purified using RP-HPLC. The synthesized **2c** or **5c** were dissolved in dry THF, the pH was adjusted to around 8 using DIEA, and the mixture was chilled in an ice-water bath and while being flushed with nitrogen. Then, 4 equivalents of the acetyl chloride were added dropwise, and the temperature was raised to room temperature. The reaction was left standing for 3 hours and then quenched with ice-cold water. After the solvent was removed by rotary evaporation, the resulting oily product was purified directly by RP-HPLC to yield **2b** or **5b** (**Scheme S3**).

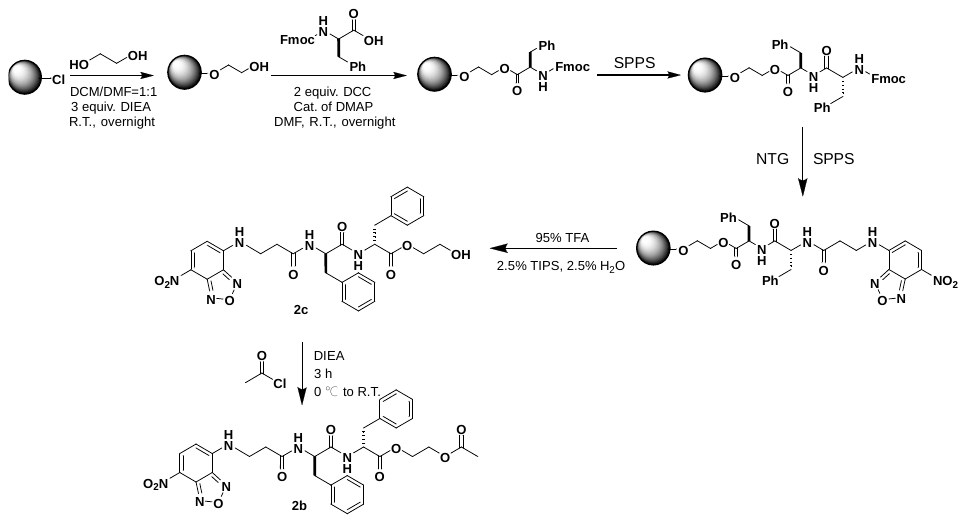

**Scheme S3.** Synthetic procedure of **2b** and **2c**.

**Synthesis of 2d**

**2d** was synthesized based on **2a** with the reported method^1^. Briefly, 0.12 mmol of **2a** was dissolved in CH_3_CN (2 mL), and H_2_O_2_ (30%, 1.2 mmol) and CH_3_ReO_3_ (0.012 mmol) were added to the solution. The reaction was stirred for 2 hours at room temperature. Afterward, the solvent is air-dried, and the remaining oily product was dissolved in methanol and purified by RP-HPLC to obtain **2d**.

**Synthesis of 5d**

Briefly, 0.1 mmol of synthesized peptide (BPff) and 3 equivalents of DIEA were dissolved in 2 mL dry DCM, followed by the dropwise addition of 5 equivalents of thionyl chloride. The reaction was stirred for 30 min, and 10 mL of methanol was added, followed by stirring for 3 hours. Afterward, the solvent is air-dried, and the remaining oily product was dissolved in methanol and purified by RP-HPLC to obtain **5d**.

**Supplementary figures**

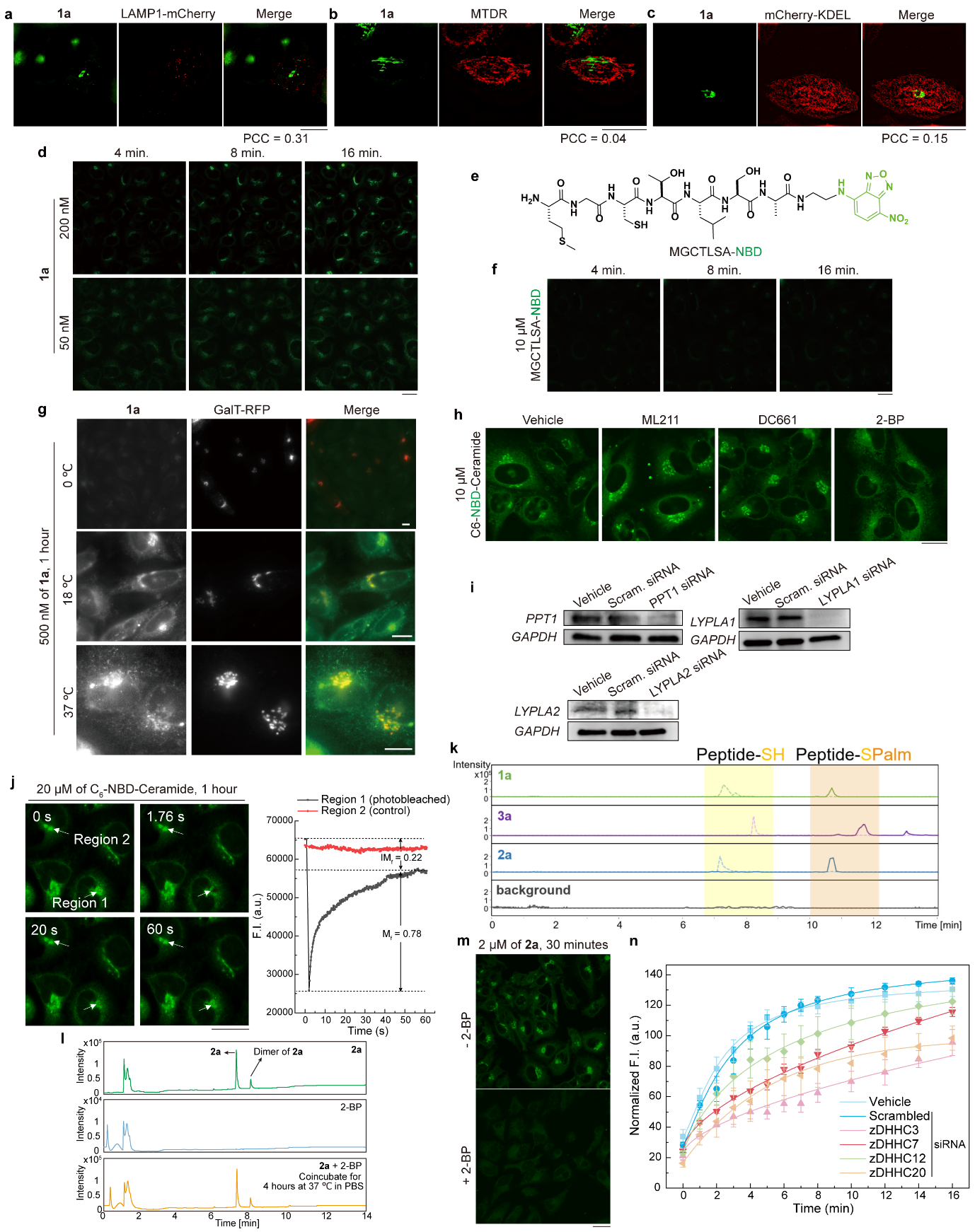

**Supplementary Figure S1.** **Non-diffusive CyMA specifically accumulate at the Golgi.** (a-c) Colocalization study of CyMA with lysosome (a), mitochondria (b), ER (c). (d) CLSM of HeLa cells treated with **1a** (200 nM, 50 nM; 4, 8, 16 min.). (e) Molecular structure of MGCTLSA-NBD as a control. (f) CLSM of HeLa cells treated with MGCTLSA-NBD (10 μM; 4, 8, 16 min.). (g) CLSM of HeLa cells treated with **1a** (500 nM; 1 hour) at different temperatures. (h) CLSM of HeLa cells pretreated with or without ML211 (50 μM; 30 min.) or DC661 (20 μM; 30 min.) or 2-BP (20 μM; 30 min.) and then treated with C6-NBD-Ceramide (20 μM; 16 min.). (i) Immunoblotting of PPT1, LYPLA1, LYPLA2 and GAPDH in HeLa cells transfected with the corresponding siRNA. (j) FRAP assay of the Golgi in HeLa cells treated with C6-NBD-Ceramide (20 μM; 1 hour). (k) Retention time comparison of the corresponding Peptide-SH and Peptide-SPalm resulting from different CyMA (**1a**, **2a**, and **3a**) using LC-HRMS. (l) LC trace of two starting materials, **2a** and 2-BP, and their mixture incubated for 4 h at 37 °C in PBS. (m) CLSM of HeLa cells pretreated with or without 2-BP (20 μM; 30 min.) and then treated with **2a** (2 μM; 30 min.). (n) Quantification of the fluorescence intensity at GA of zDHHC siRNA transfected HeLa cells treated with **1a** (500 nM) within 16 minutes. Scale bar = 20 μm except for (g), where scale bar = 10 μm.

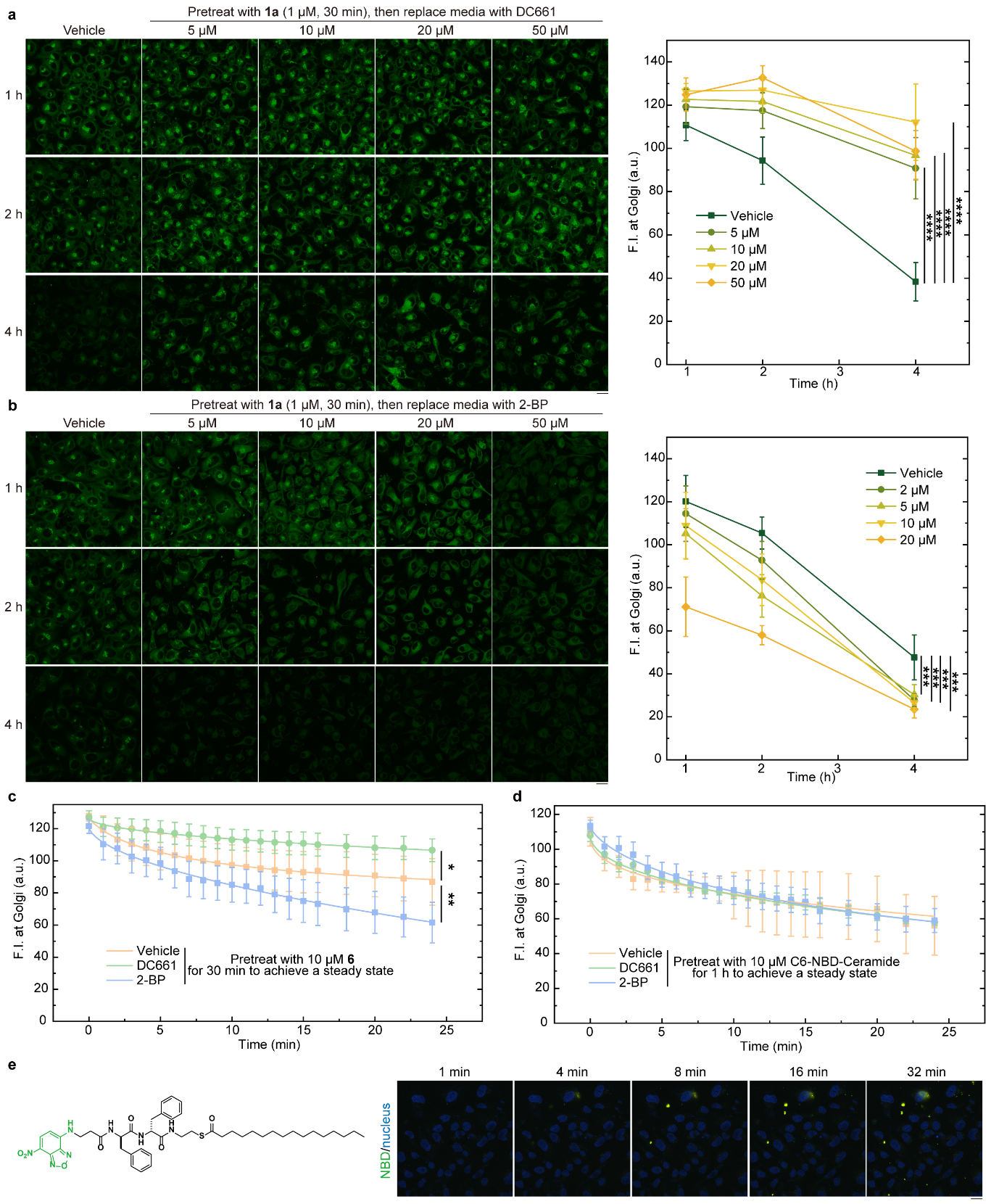

**Supplementary Figure S2.** **CyMA establish futile cycles by targeting an enzyme switch.** (a) CLSM of HeLa cells pretreated with **1a** (1 μM; 30 min.), followed by replacement of the media with varying concentrations of PPT1 inhibitor, DC661, for 1, 2, 4 hours; Quantification of fluorescence intensity at the Golgi is shown. (b) CLSM of HeLa cells pretreated with **1a** (1 μM; 30 min.), followed by replacement of the media with varying concentrations of zDHHC inhibitor, 2-BP, for 1, 2, 4 hours; Quantification of fluorescence intensity at the Golgi is shown. (c) Quantitative analysis of the fluorescent intensity at the Golgi in HeLa cells pretreated with **6** (10 μM, 30 min.), followed by replacement of the media with either PPT1 inhibitor (DC661, 20 μM) or zDHHC inhibitor (2-BP, 20 μM). (d) Quantitative analysis of the fluorescent intensity at the Golgi of HeLa cells pretreated with C6-NBD-Ceramide (10 μM, 1 hour), followed by replacement of the media with either PPT1 inhibitor (DC661, 20 μM) or zDHHC inhibitor (2-BP, 20 μM). (e) Molecular structure of the palmitoylated CyMA and the CLSM of its co-incubation (10 μM) with HeLa cells for 1, 4, 8, 16, 32 minutes. Scale bar = 20 μm.

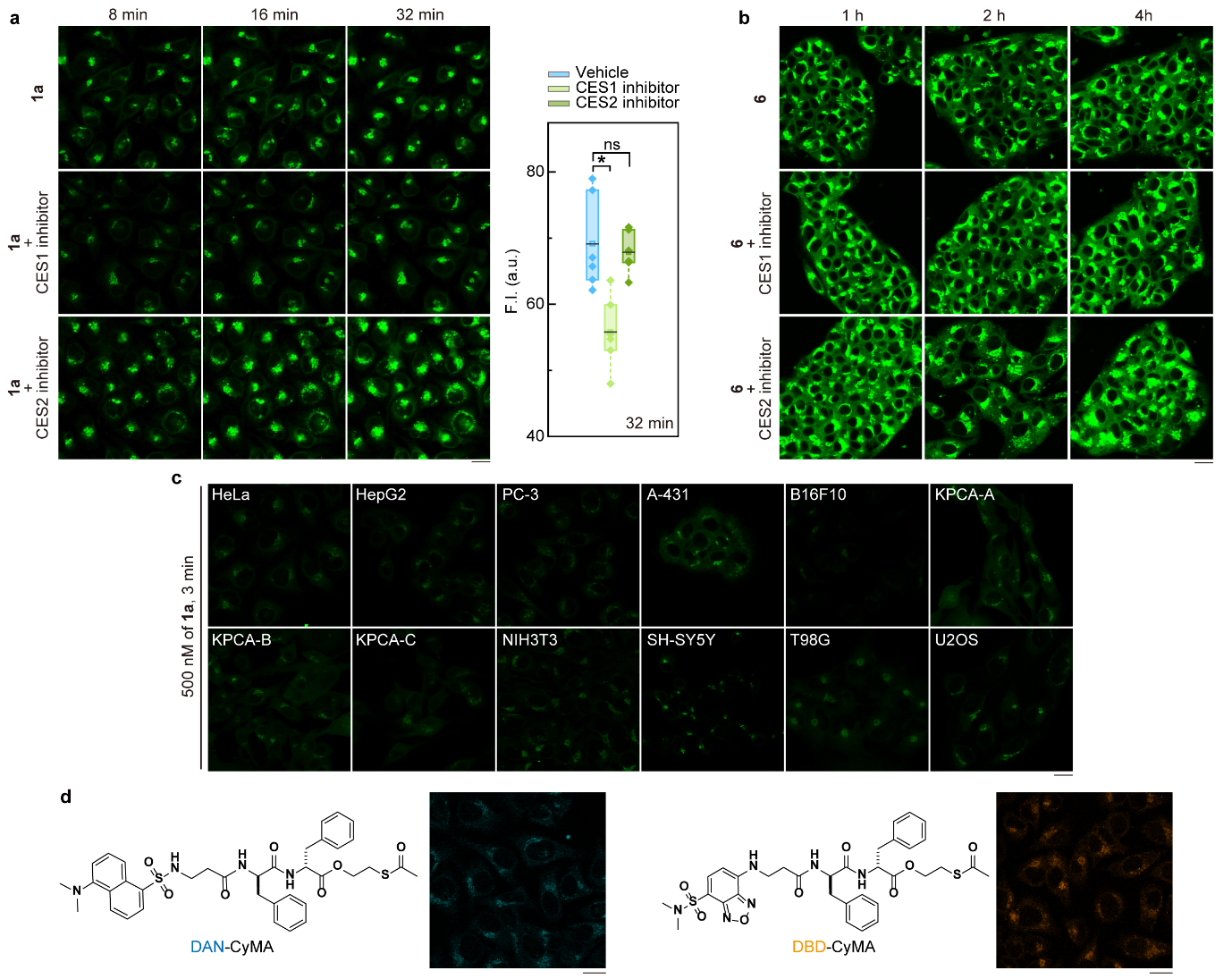

**Supplementary Figure S3.** **CES-responsive CyMA as versatile GA probes.** (a) CLSM of HeLa cells pretreated with vehicle, CES1 inhibitor (Nevadensin, 20 μM) or CES2 inhibitor (Loperamide, 20 μM) for 30 min., followed by co-treatment with **1a** (1 μM, 8, 16, 32 min.). (b) CLSM of HepG2 cells pretreated with vehicle, CES1 inhibitor (Nevadensin, 20 μM) or CES2 inhibitor (Loperamide, 20 μM) for 30 min., followed by co-treatment with **6** (5 μM, 1, 2, 4 hour). (c) CLSM of various cell lines (HeLa, HepG2, PC-3, A-431, B16F10, KPCA-A, KPCA-B, KPCA-C, NIH3T3, SH-SY5Y, T98G, U2OS) treated with **1a** (500 nM, 3 min.). (d) Molecular structures of DAN- and DBD-conjugated CyMA, with corresponding CLSM images of HeLa cells treated with DAN-CyMA (500 nM, 30 min.) or DBD-CyMA (5 μM, 15 min.). Scale bar = 20 μm.

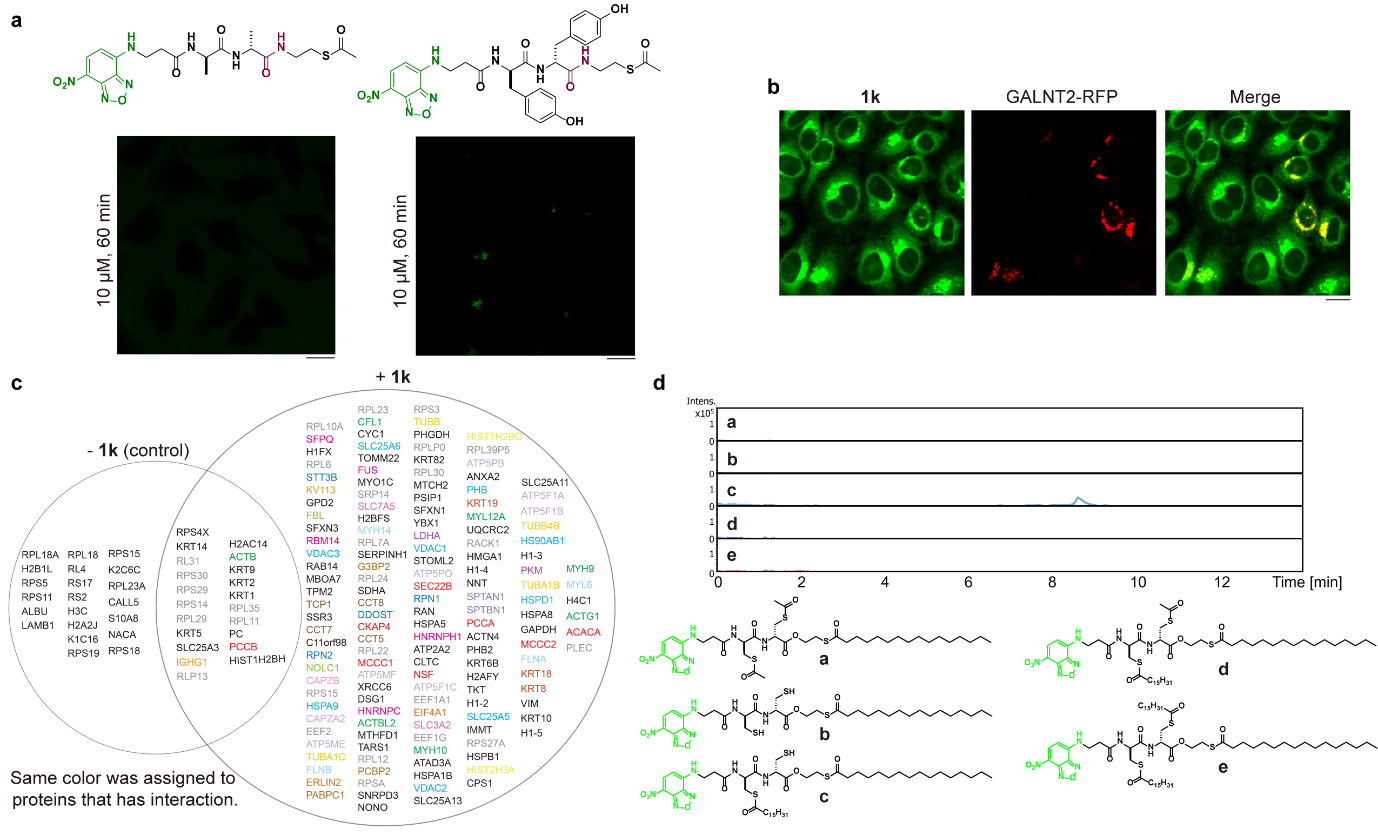

**Supplementary Figure S4.** **Generality of CyMA targeting the Golgi without direct protein binding.** (a) CLSM of HeLa cells treated with analogues **1e** and **1f** (10 μM, 60 min.), in which ester bonds are replaced with amide bonds. (b) CLSM of GALNT2-RFP transfected HeLa cells treated with **1k** (5 μM, 30 min.). (c) Proteins pulled down via UV-crosslinking of **1k** analyzed by LC-MS/MS. (d) Extracted-ion chromatogram (EIC) in LC-HRMS of potential components present in the lysates of HeLa treated with **1l** (2 μM, 30 min.). Scale bar = 20 μm.

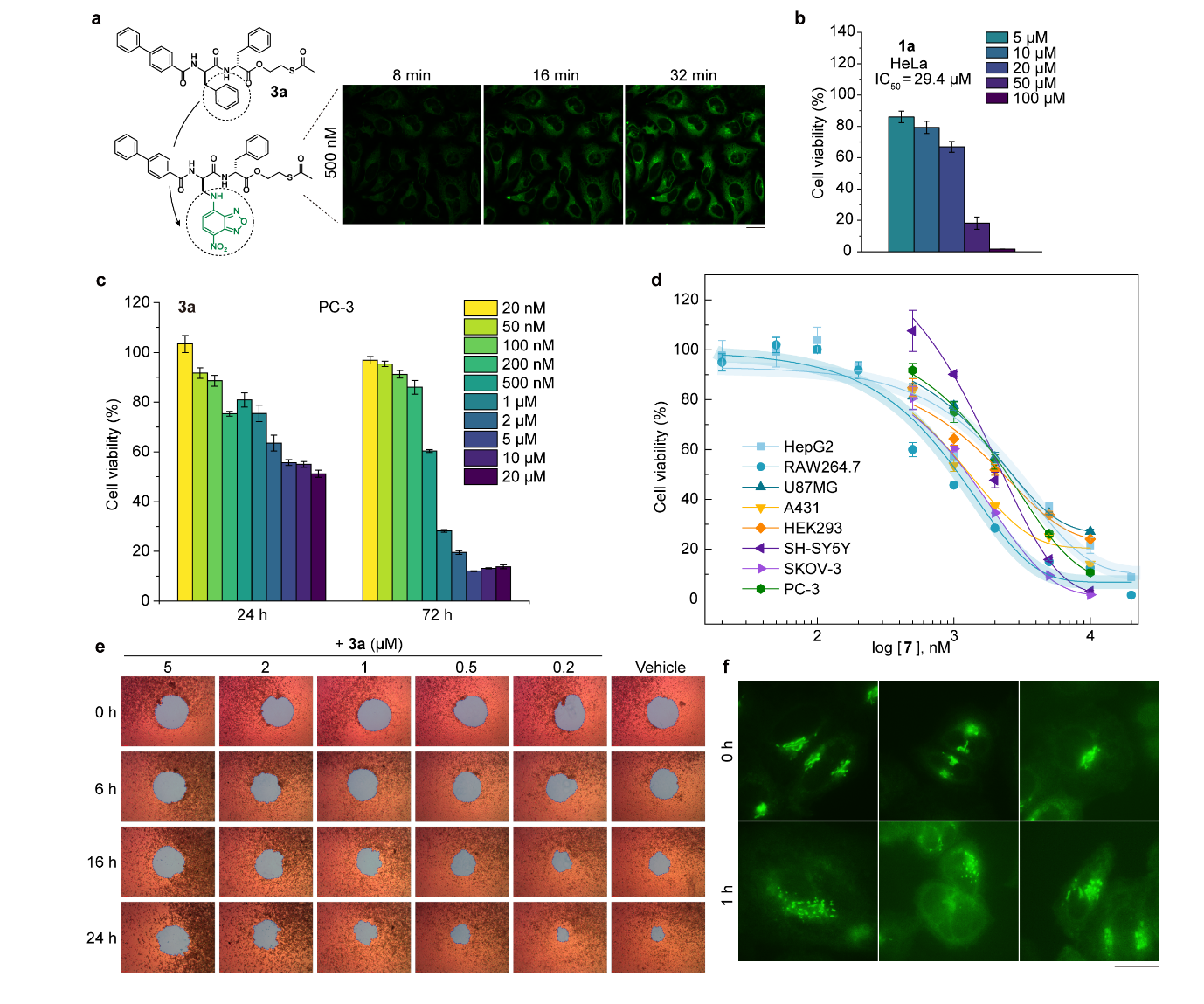

**Supplementary Figure S5.** **CyMA inhibit proliferation and migration of cancer cells with high efficacy by disrupting Golgi.** (a) CLSM of HeLa cells treated with an analogue of **3a** (500 nM), in which the benzene ring is replaced with a hydrophobic fluorophore for 8, 16, 32 minutes. (b) Cell viability of HeLa cells treated with **1a** for 24 hours. (c) Cell viability of PC-3 cells treated with **3a** for 24 hours and 72 hours. (d) Cell viability of various cells with different CES expression levels treated with **7** for 24 h; Two cell lines (HepG2 and RAW264.7) overexpressing CES are highlighted. (e) Light microscope images of 2D-migration performance of HeLa cells treated with or without **3a**. (f) Three representative CLSM images of HeLa cells treated with **3a** (5 μM, 1 hour) combined with **1a** (2 μM, 1 hour). Scale bar = 20 μm.

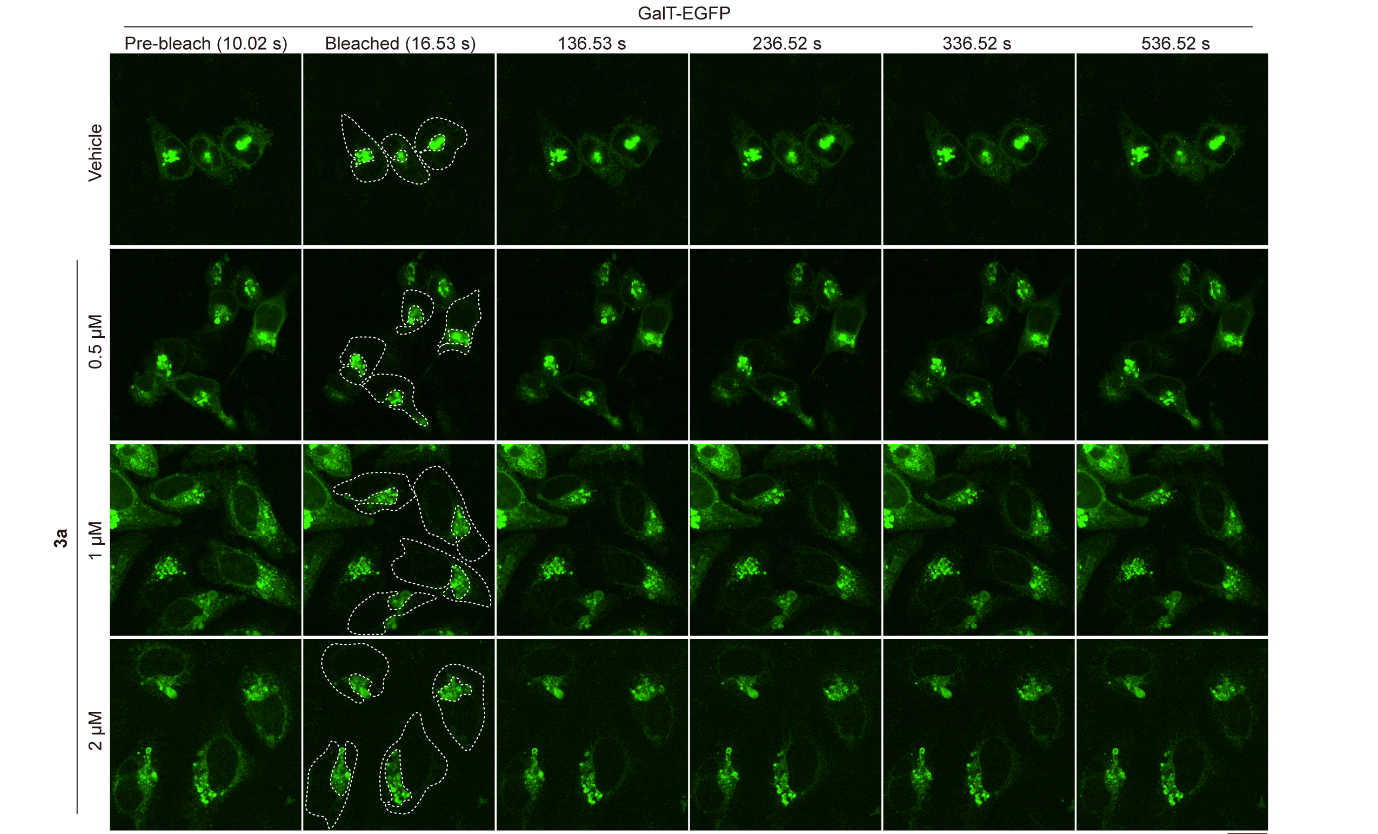

**Supplementary Figure S6.** Photobleaching of the ER pool of GalT-EGFP in HeLa cells treated with **3a** (500 nM, 1 μM, 2 μM) for 6 hours. The regions of interest are marked. Scale bar = 20 μm.

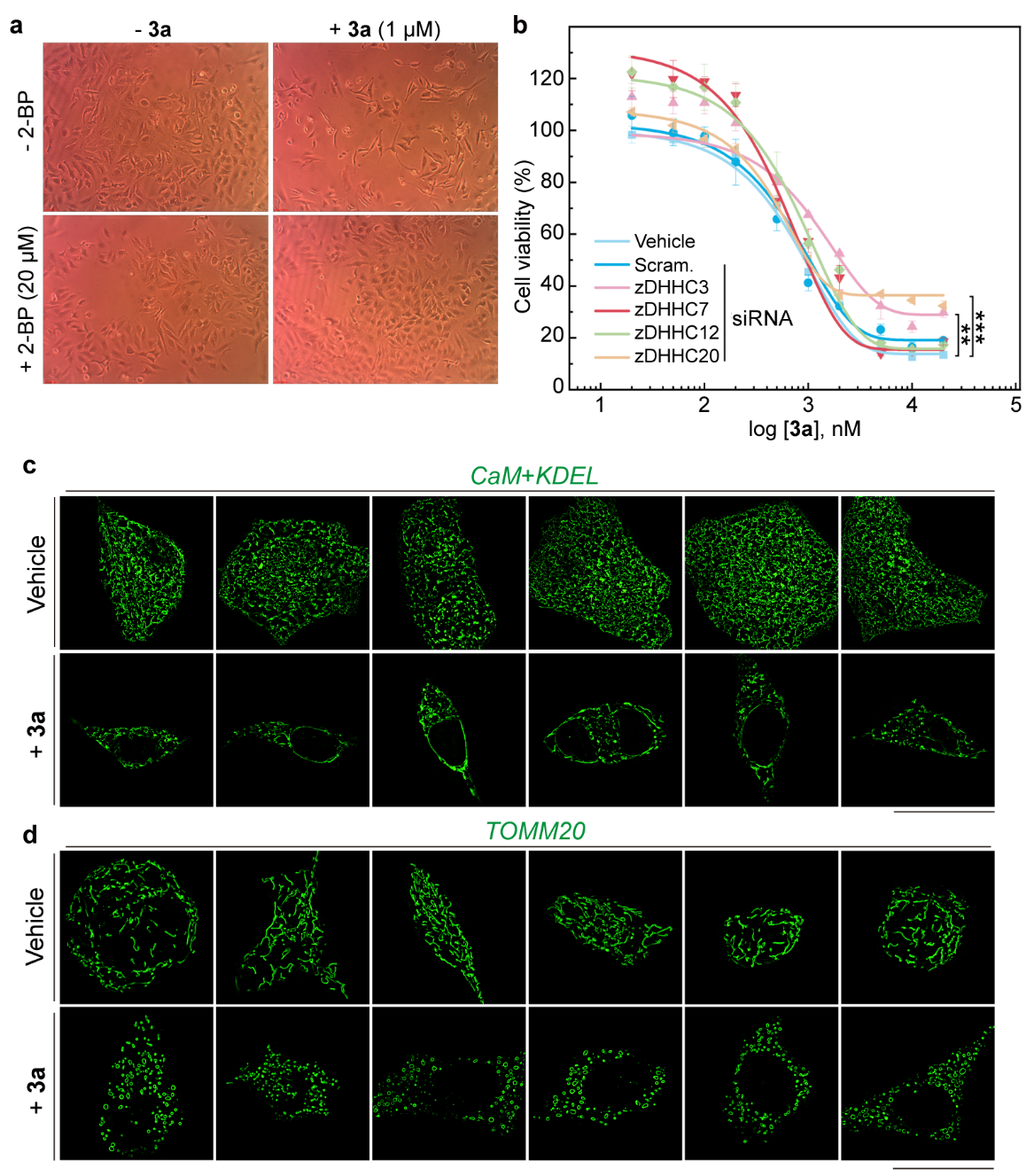

**Supplementary Figure S7.** **Inhibition or knockdown of zDHHCs rescues cells from CyMA that disrupt the Golgi, ER, and mitochondria.** (a) Light microscope images of HeLa cells treated with or without **3a** (1 μM) and 2-BP (20 μM) for 24 hours. (b) Cell viability of zDHHC siRNA transfected HeLa cells treated with **3a** for 24 hours. (c) SIM images of the ER in HeLa cells treated with or without **3a** (2 μM, 24 hours). (d) SIM images of mitochondria in HeLa cells treated with or without **3a** (2 μM, 24 hours). Scale bar = 20 μm.

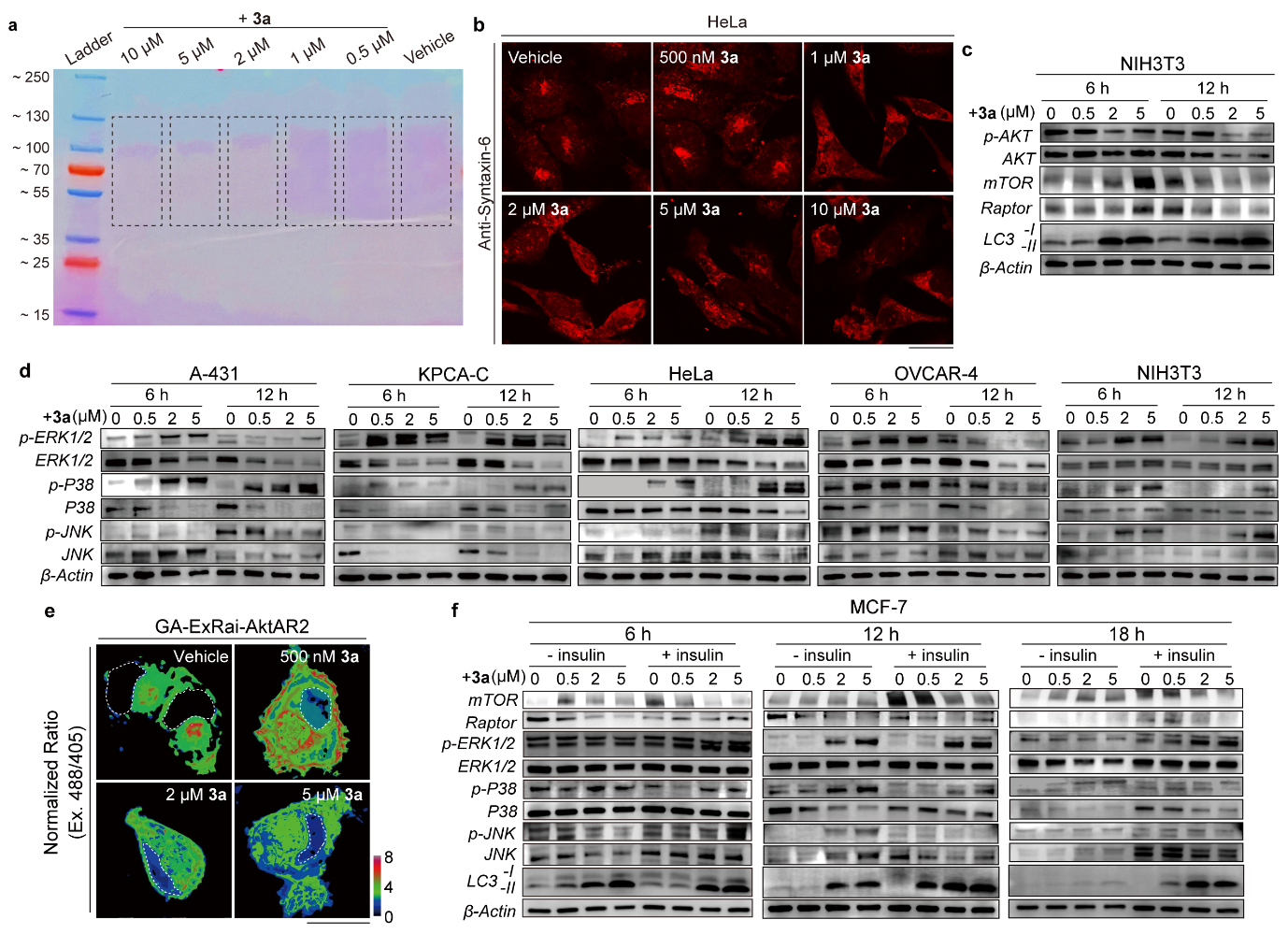

**Supplementary Figure S8.** **Mechanistic studies of CyMA that disrupt signaling pathways leading to cell death.** (a) Glycoprotein staining of SDS-PAGE of the whole cell lysate from KPCA-C cells (10 μg) treated with **3a** (10, 5, 2, 1, 0.5 μM) or vehicle for 6 hours. (b) Immunocytochemistry of Syntaxin-6 in HeLa cells treated with or without **3a** (500 nM, 1 μM, 2 μM, 5 μM, 10 μM; 6 hours). (c) Immunoblotting of key proteins (p-AKT, AKT, mTOR, Raptor, LC3B) in the AKT-mTOR signaling pathway in NIH3T3 cells treated with or without **3a** (500 nM, 2 μM, 5 μM; 6 and 12 hours). (d) Immunoblotting of key proteins (p-ERK, ERK, p-P38, P38, p-JNK, JNK) in the Ras/Raf/MAPK signaling pathway across different cell lines treated with or without **3a** (500 nM, 2 μM, 5 μM; 6 and 12 hours). (e) Determination of AKT activity using a fluorescent biosensor, GA-ExRai-AktAR2. (f) Immunoblotting of key proteins (mTOR, Raptor, LC3B) in the AKT-mTOR signaling pathway and key proteins (p-ERK, ERK, p-P38, P38, p-JNK, JNK) in the Ras/Raf/MAPK signaling pathway in MCF-7 cells stimulated with or without insulin and treated with or without **3a** (500 nM, 2 μM, 5 μM; 6, 12 and 18 hours).

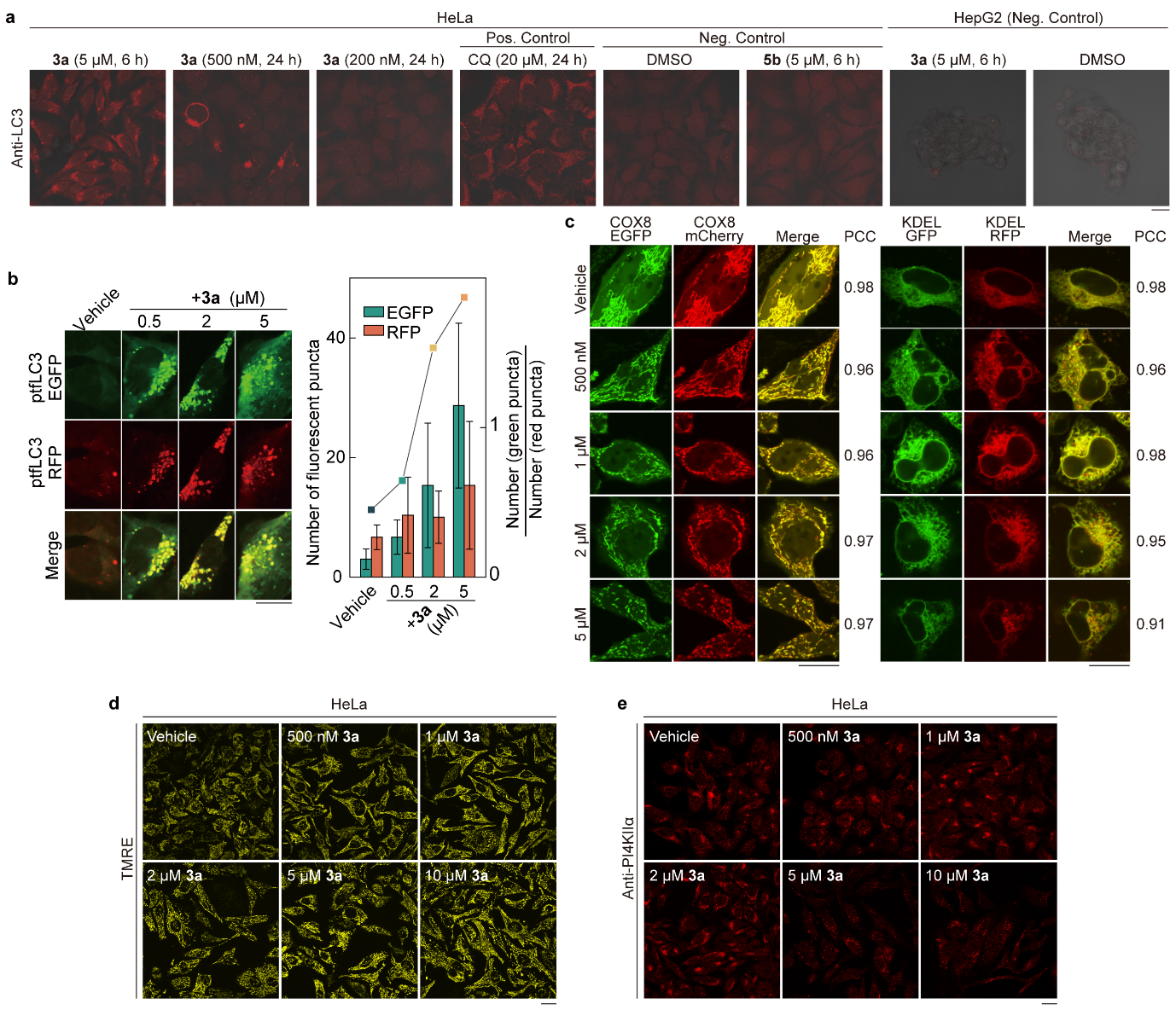

**Supplementary Figure S9.** **CyMA promote autophagosome formation but inhibit its further maturation.** (a) Immunocytochemistry of LC3 in HeLa cells and HepG2 cells under varying conditions as described; CQ represents chloroquine. (b) CLSM of ptfLC3-HeLa cells treated with or without **3a** (500 nM, 2 μM, 5 μM; 6 hours), along with the quantification of green and red fluorescent puncta per cell and the ratio of green to red puncta. (c) CLSM of COX8-EGFP-mCherry-HeLa cells and KDEL-GFP-RFP-HeLa cells treated with or without **3a** (500 nM, 1 μM, 2 μM, 5 μM; 6 hours) and the corresponding Pearson correlation coefficient. (d) TMRE-mitochondrial membrane potential assay of HeLa cells treated with or without **3a** (500 nM, 1 μM, 2 μM, 5 μM, 10 μM; 6 hours). (e) Immunocytochemistry of PI3KIIα in HeLa cells treated with or without **3a** (500 nM, 1 μM, 2 μM, 5 μM, 10 μM; 6 hours). Scale bar = 20 μm.

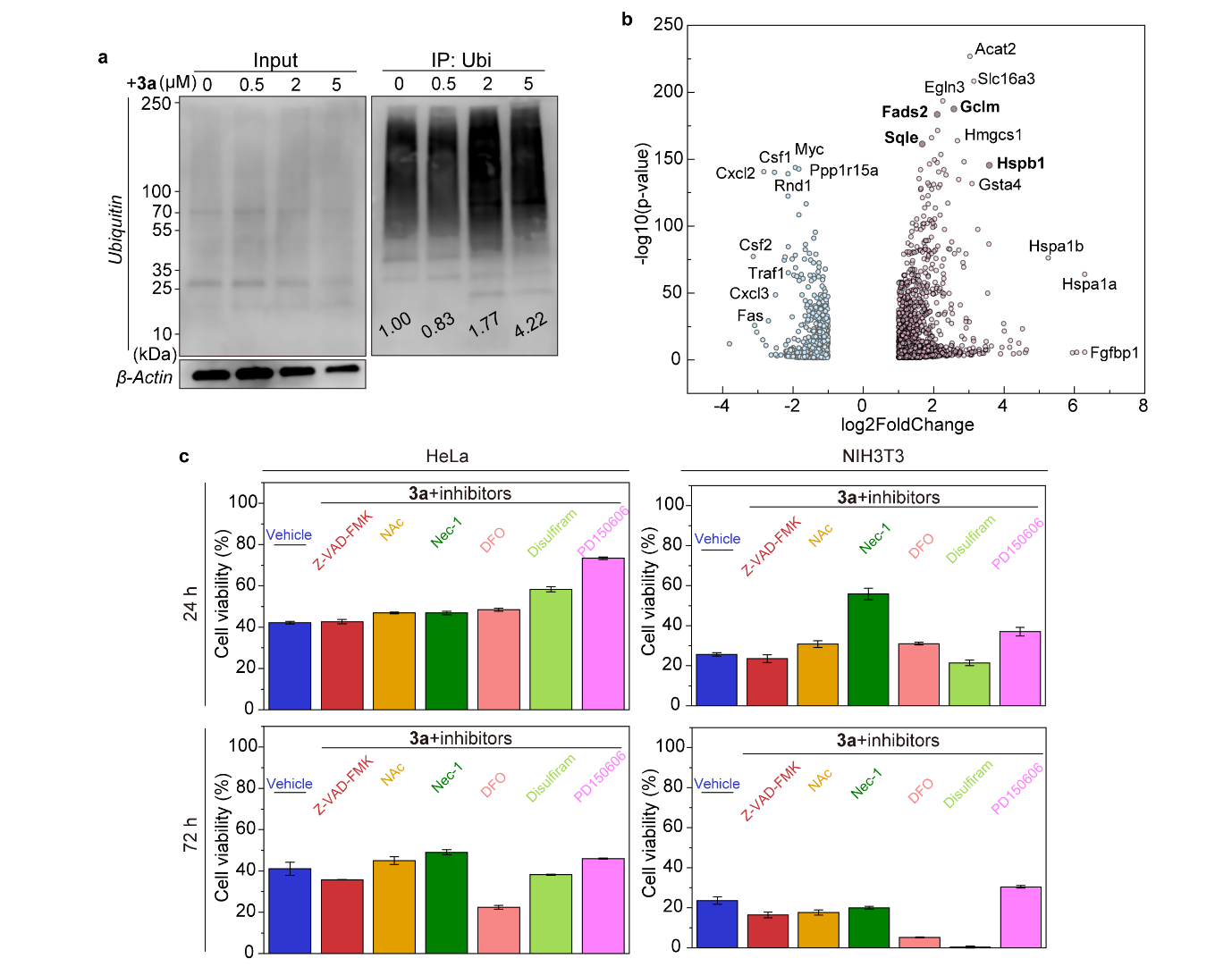

**Supplementary Figure S10.** **Multiple cell death pathways involved in CyMA-induced cell death.** (a) Immunoprecipitation of ubiquitinated proteins in lysates of KPCA-C cells treated with or without **3a** (500 nM, 2 μM, 5 μM, 6 hours) and the quantitative analysis normalized to β-Actin. (b) Volcano plot of regulated mRNA transcription in KPCA-C cells treated with **3a** (2 μM, 6 hours) identified by RNA-Seq. (c) Cell viability of HeLa and NIH3T3 cells treated with **3a** (1 μM) combined with or without cell death inhibitors (Z-VAD-FMK, 50 μM; NAc, 1mM; Nec-1, 50 μM; DFO, 5 μM; Disulfiram, 10 μM; PD150606, 100 μM) for 24 and 72 hours.

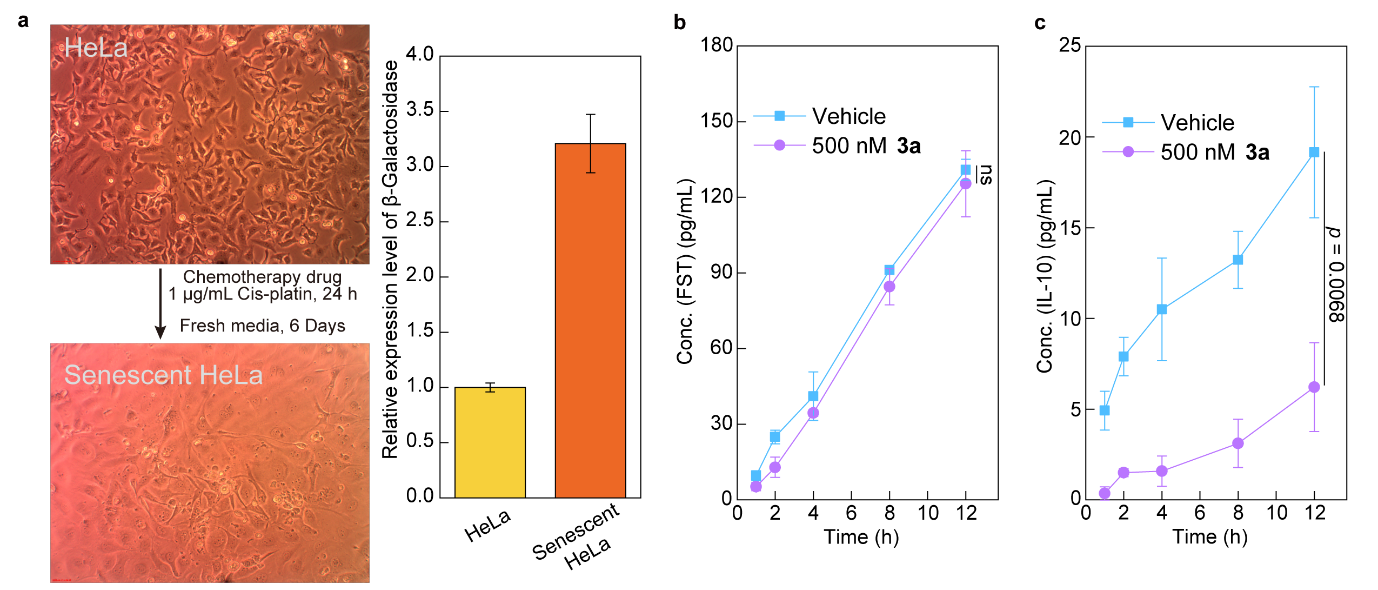

**Supplementary Figure S11.** **CyMA modulate TME by inhibiting secretion of pro-cancer cytokines.** (a) Light microscope images of HeLa cells and senescent HeLa cells, evident by the difference in β-Galactosidase expression level between the two. (b) ELISA of FST in the conditioned media of KPCA-C after the treatment of vehicle or **3a** (500 nM). (c) ELISA of IL-10 in the conditioned media of KPCA-C after the treatment of vehicle or **3a** (500 nM).

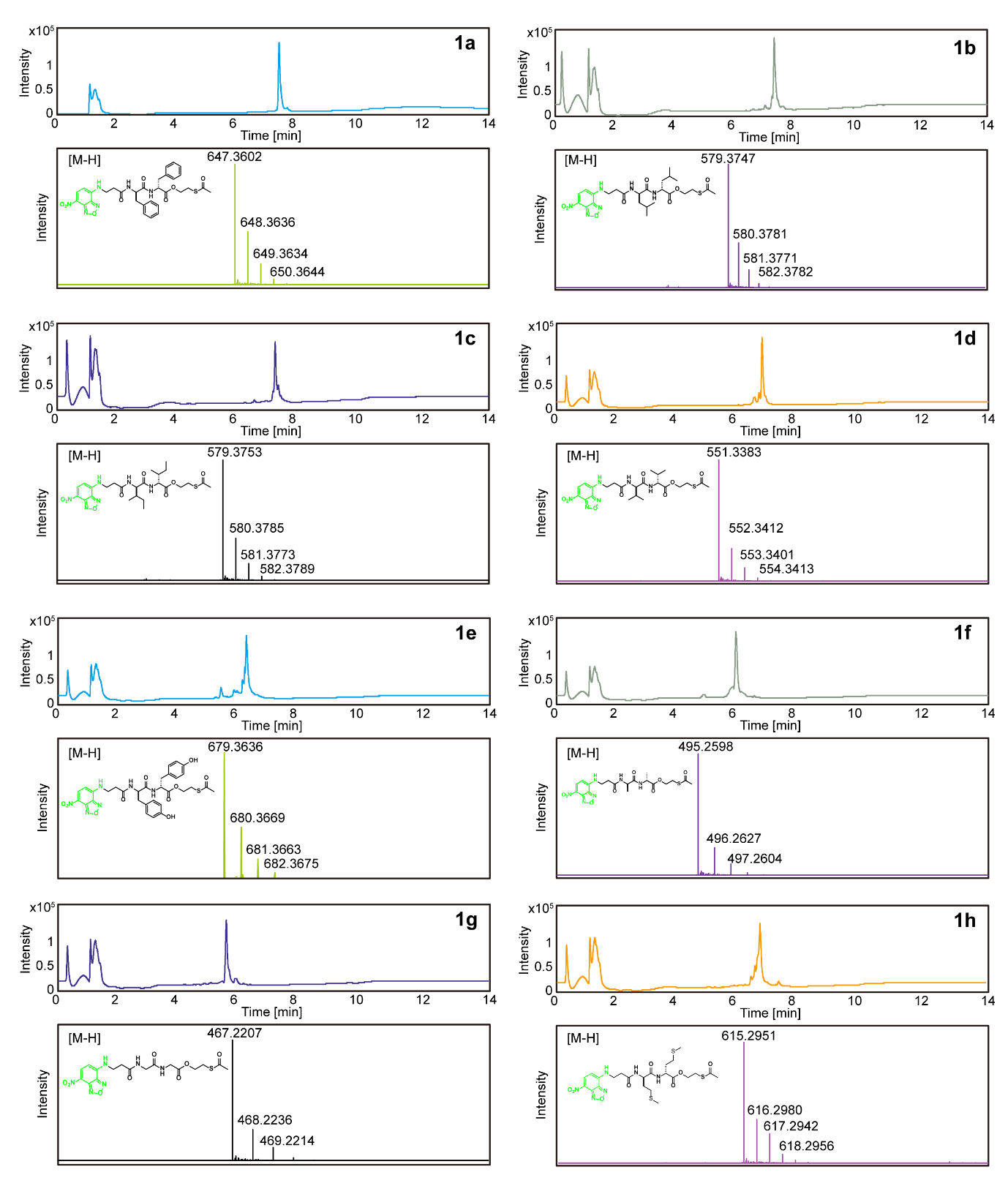

**Supplementary Figure S12.** LC-HRMS of the synthesized **1a**-**1h**.

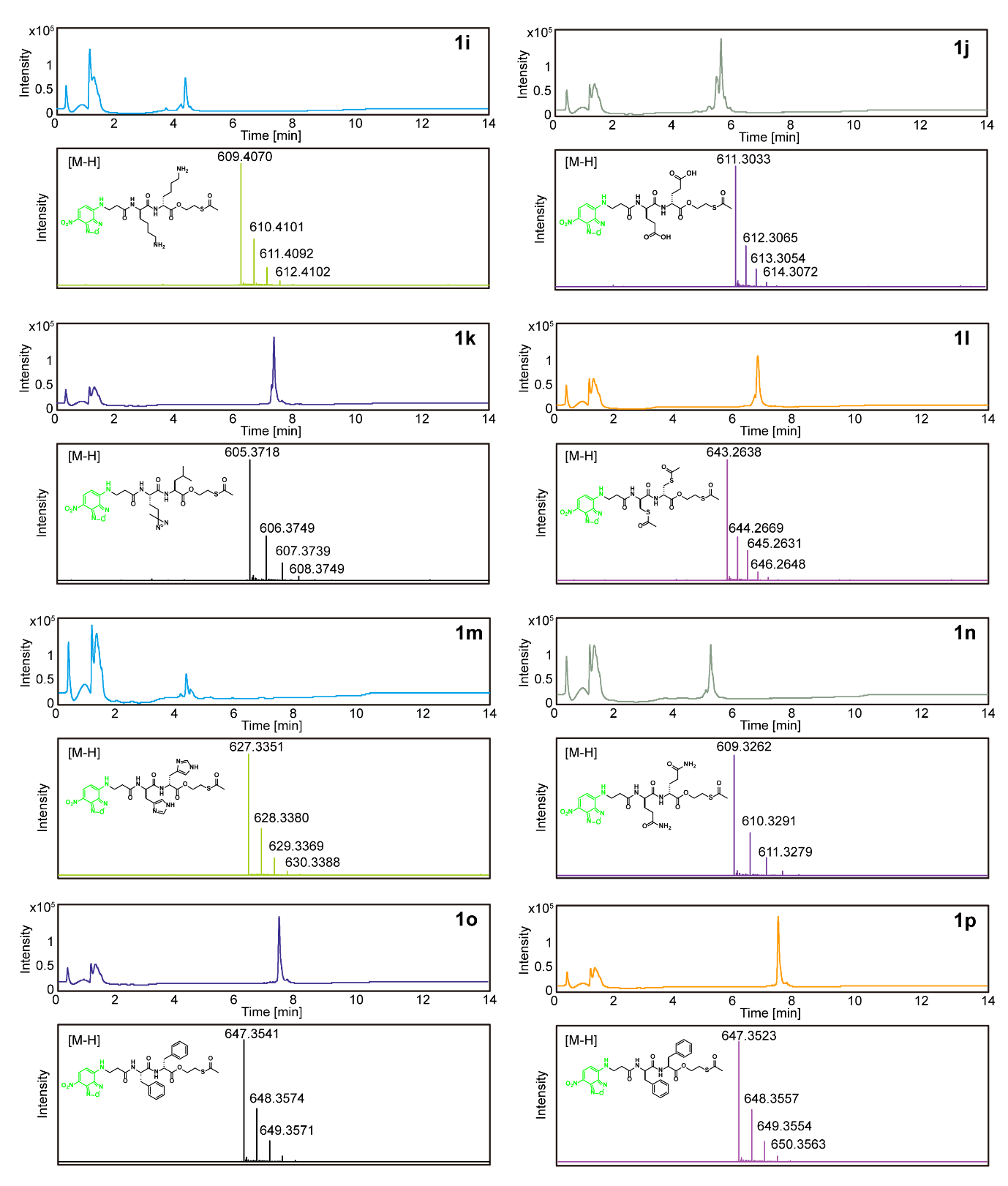

**Supplementary Figure S13.** LC-HRMS of the synthesized **1i**-**1p**.

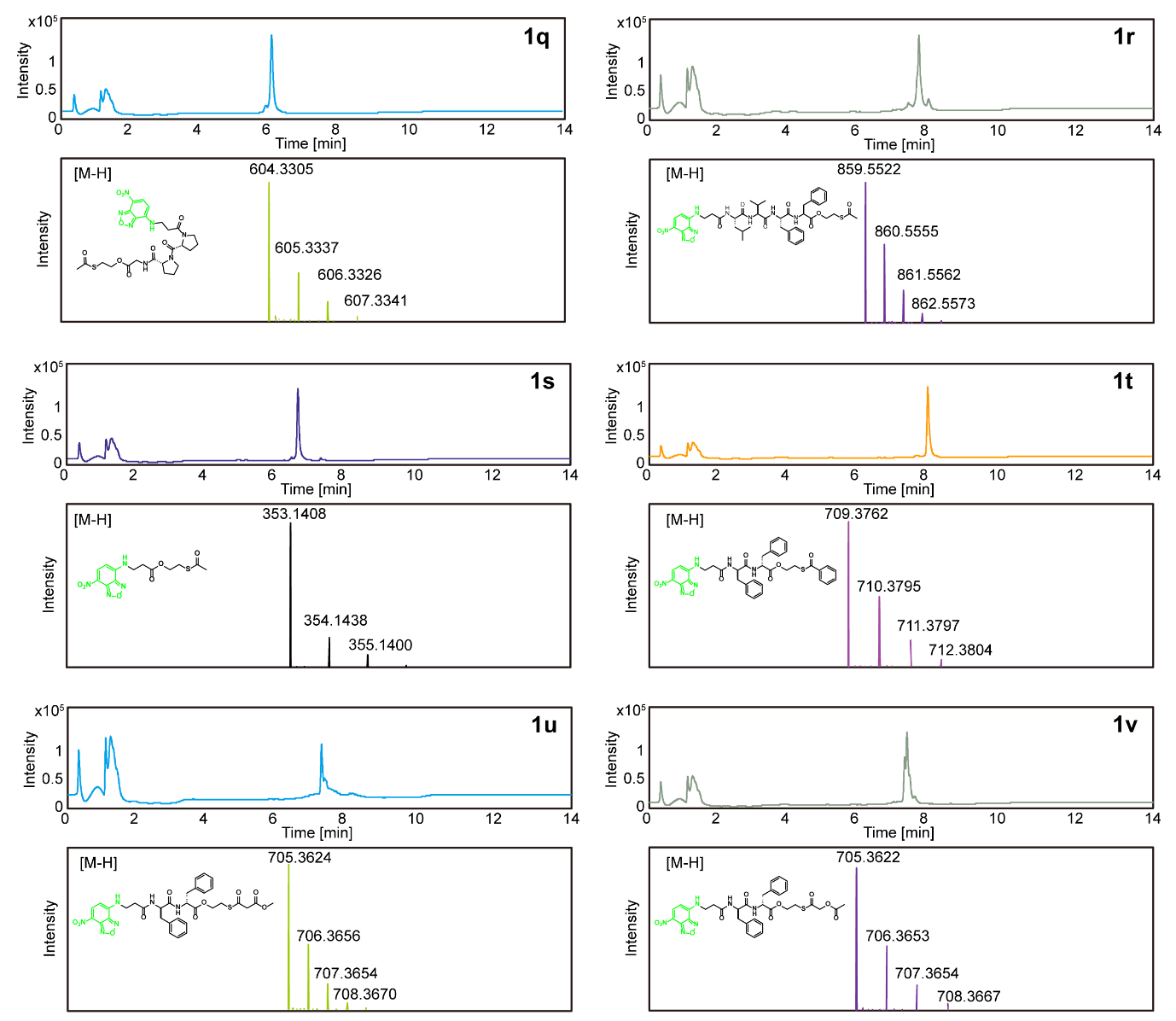

**Supplementary Figure S14.** LC-HRMS of the synthesized **1q**-**1v**.

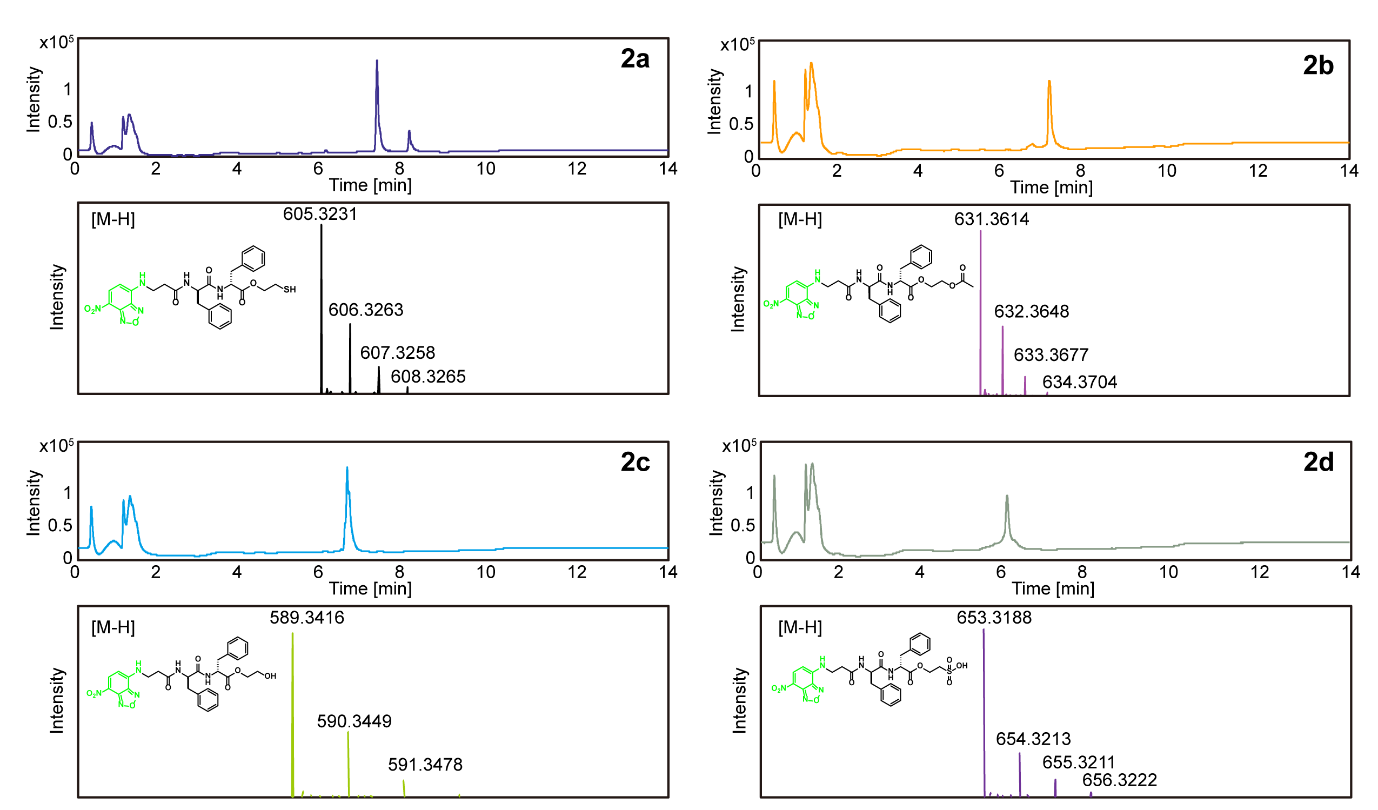

**Supplementary Figure S15.** LC-HRMS of the synthesized **2a**-**2d**.

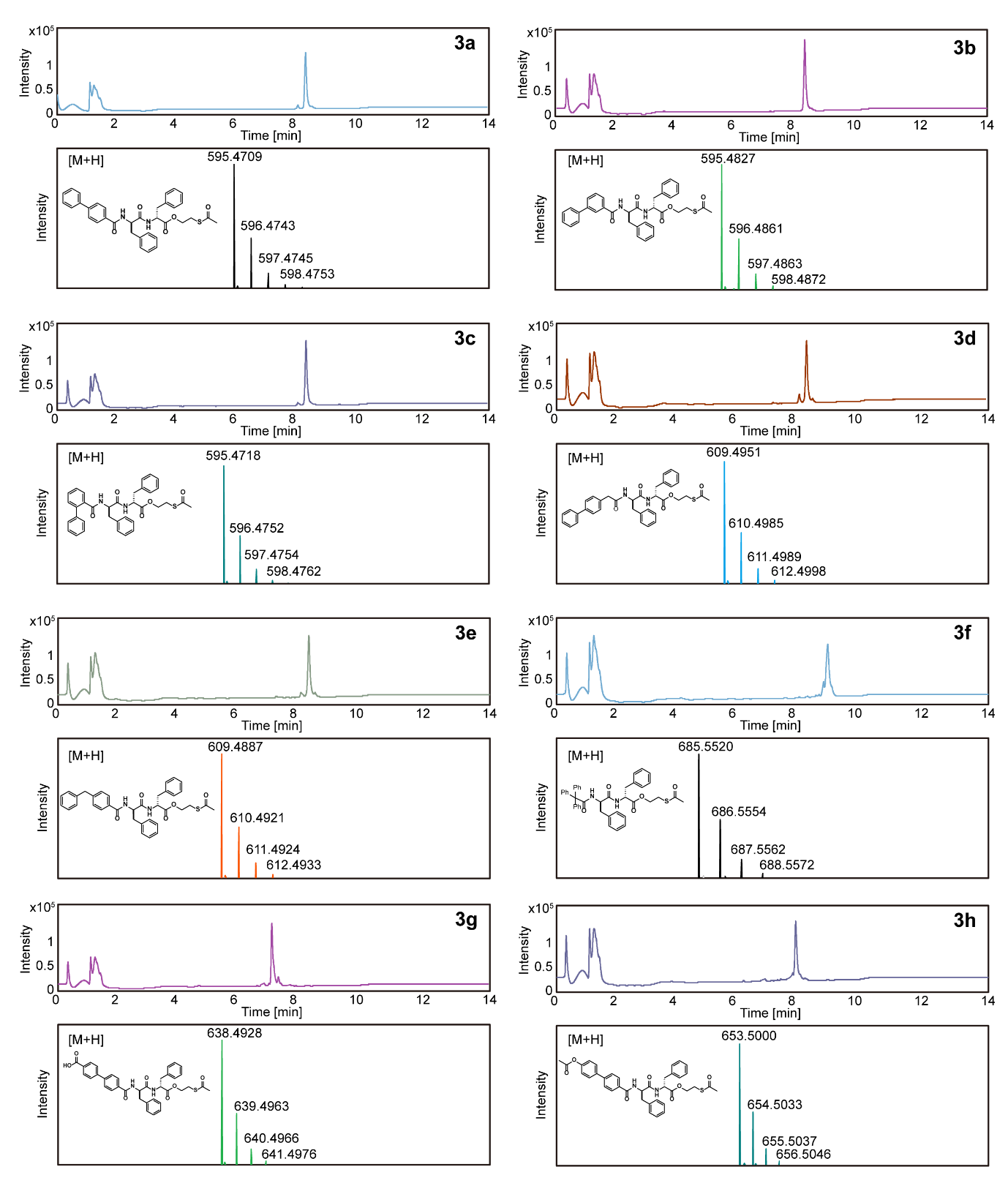

**Supplementary Figure S16.** LC-HRMS of the synthesized **3a**-**3h**.

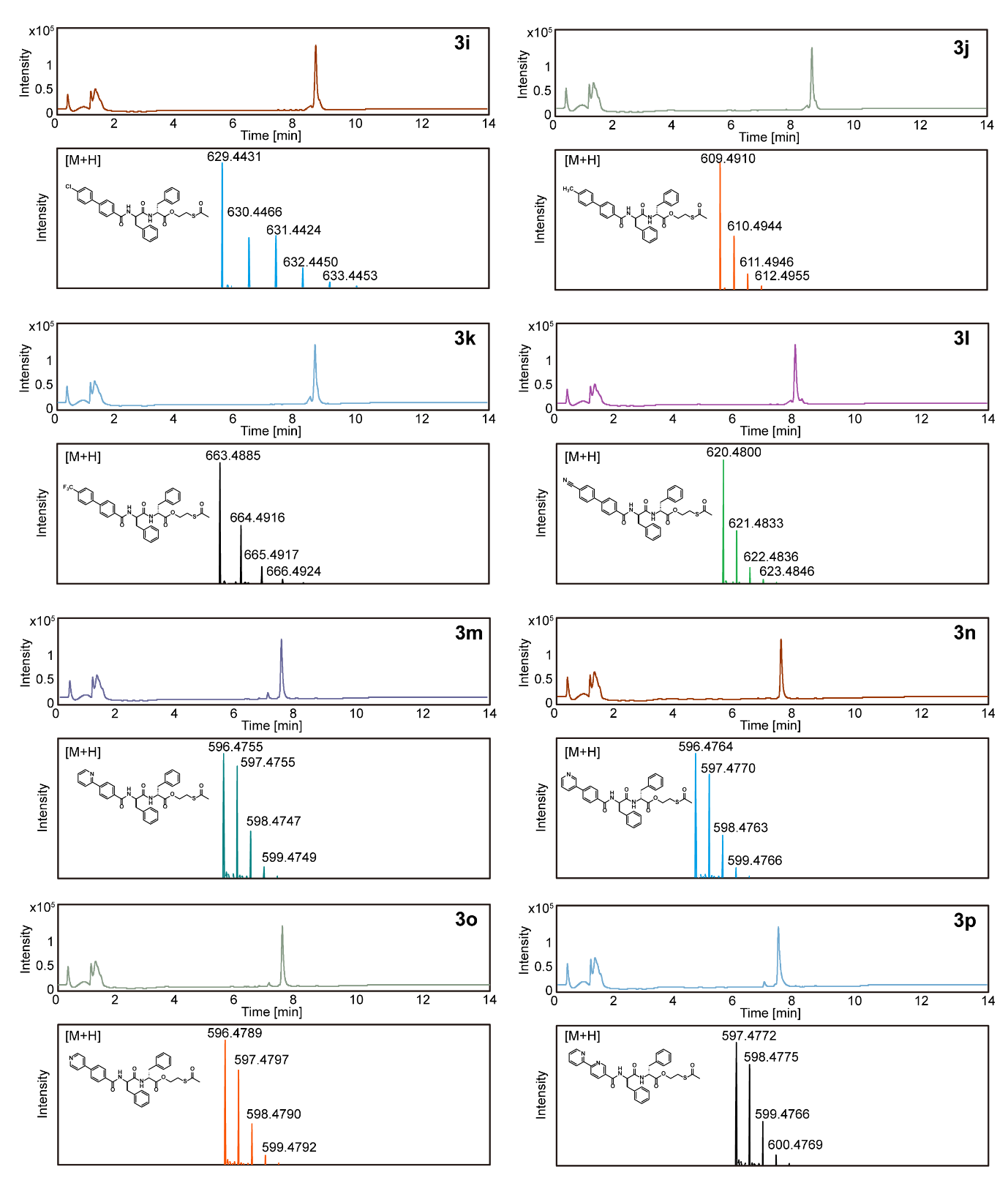

**Supplementary Figure S17.** LC-HRMS of the synthesized **3i**-**3p**.

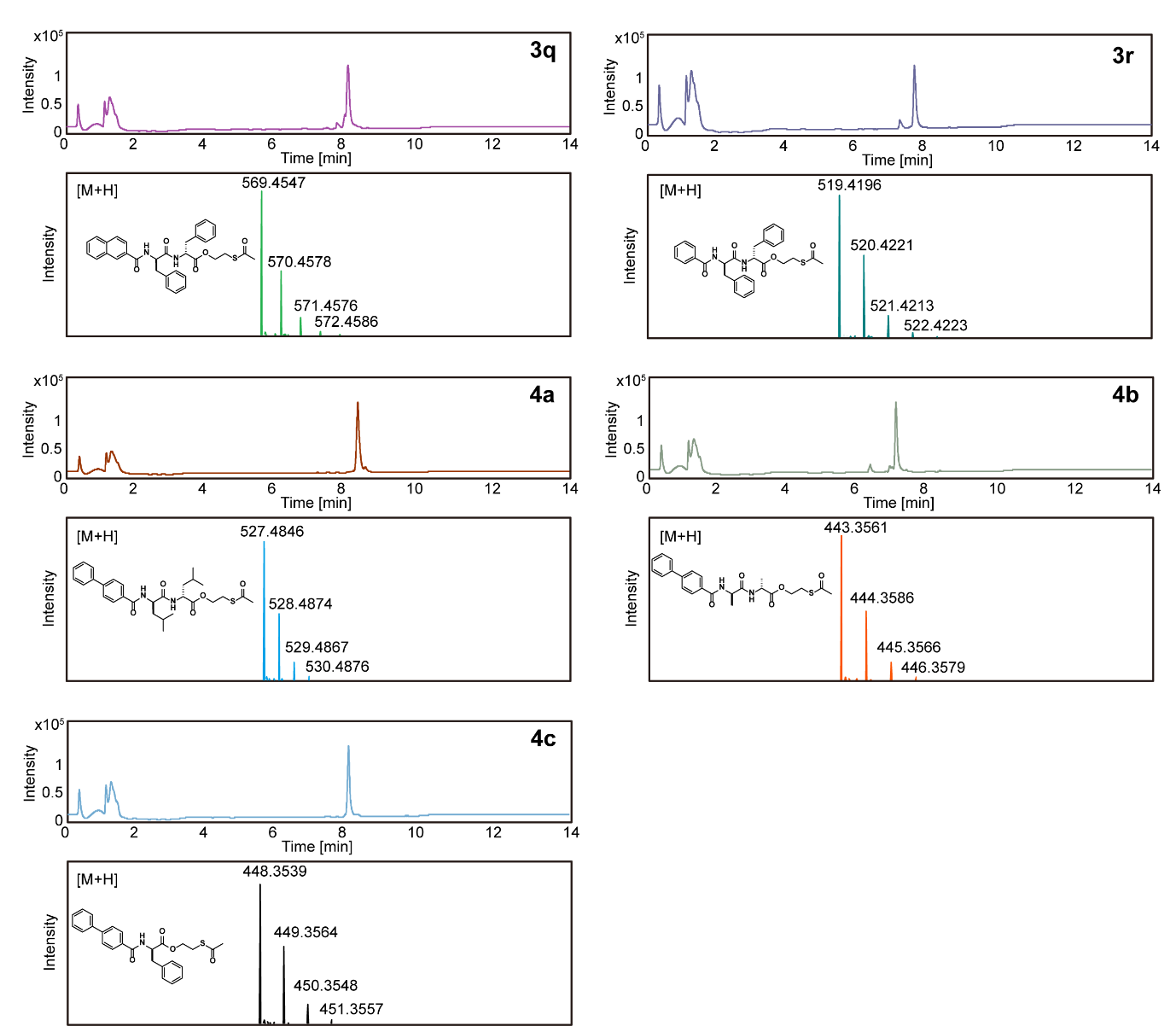

**Supplementary Figure S18.** LC-HRMS of the synthesized **3q**-**3r**, **4a**-**4c**.

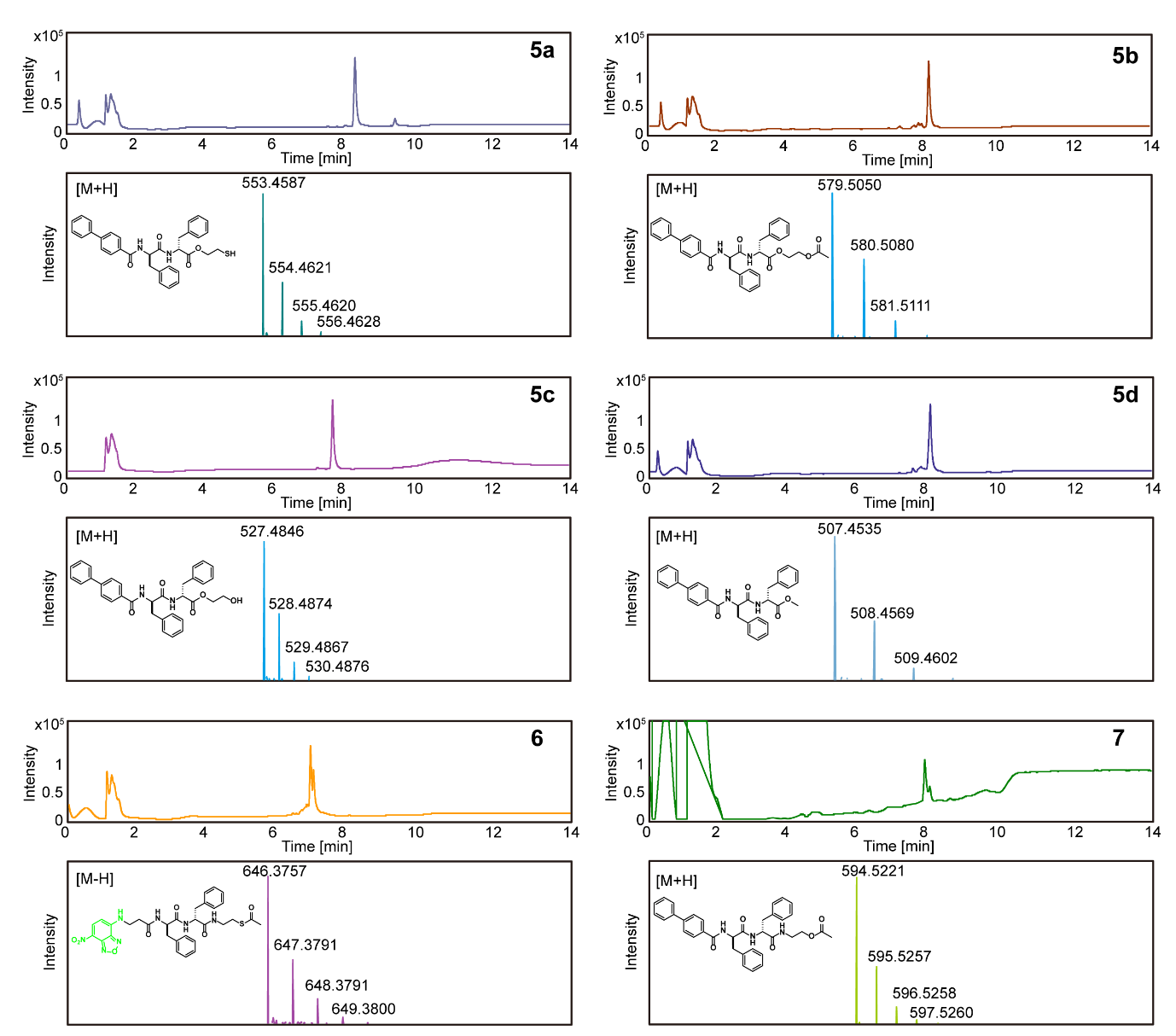

**Supplementary Figure S19.** LC-HRMS of the synthesized **5a**-**5d**, **6**, **7**.
